# A potential marker for problematic mother-infant bonding revealed by magnetoencephalography study of first-time mothers listening to infant cries

**DOI:** 10.1101/2022.06.24.497467

**Authors:** N.F Hoegholt, L. Bonetti, A.B.A. Stevner, C.E. Andersen, M Hughes, H.M. Fernandes, P. Vuust, M.L Kringelbach

**Affiliations:** Center for Music in the Brain, Department of Clinical Medicine, Aarhus University & The Royal Academy of Music Aarhus/Aalborg, 8000, Denmark; Centre for Eudaimonia and Human Flourishing, Linacre College, University of Oxford, UK; Department of Psychiatry, University of Oxford, OX37JX, Oxford, United Kingdom; Center of Functionally Integrative Neuroscience, Department of Clinical Medicine, Aarhus University, 8000, Denmark

**Keywords:** Brain, cry perception, Mother-infant relationship, parenting, sleep

## Abstract

Studies using magnetoencephalography (MEG) have identified the orbitofrontal cortex (OFC) to be an important early hub for a ‘parental instinct’ in the brain. This complements the finding from functional magnetic resonance imaging (fMRI) studies linking reward, emotion regulation, empathy and mentalisation networks to the ‘parental brain’. Here, we used MEG in 43 first-time mothers listening to infant and adult cry vocalisations to investigate the link with mother-infant postpartum bonding scores and their level of sleep deprivation (assessed using both actigraphy and sleep logs). We found significant differences 800-1000ms after onset of infant compared to adult cries in source-reconstructed brain activity in areas previously linked to the parental brain. Importantly, mothers with weaker bonding scores showed decreased brain responses to infant cries in the auditory cortex, middle and superior temporal gyrus, OFC, hippocampal areas, supramarginal gyrus and inferior frontal gyrus at around 100-200ms after stimulus onset. In contrast, we did not find correlations with sleep deprivation scores. The significant changes in brain processing of an infant’s distress signals could be a novel marker of weaker infant bonding in new mothers and should be investigated in vulnerable populations.

---

Becoming a parent for the first time is a life-changing event that has given rise to significant scientific interest. While several studies in the field originally focused on the evolutionary aspects of parent-infant communication (Darwin, 1872, 1877; Lorenz, 1943), more recent lines of research have focused on the attachment relationships and the maternal sensitivity (Ainsworth, 1978; Bowlby, 1997). With the introduction of neuroimaging methods such as electroencephalography (EEG), functional magnetic resonance imaging (fMRI), and magnetoencephalography (MEG), new insights into the ‘parental brain’ have emerged. Throughout the latest years, several models of the ‘parental brain’ have been suggested (Feldman, 2015; Fuths et al., 2017; Kim et al., 2016; Kringelbach et al., 2016; Parsons et al., 2010; Stark et al., 2019; Swain et al., 2014). The models are based on different neuroimaging methods but all models involve brain networks previously identified as involved in reward, emotion regulation, empathy and mentalisation (Feldman, 2015; Fuths et al., 2017; Kim et al., 2016; Kringelbach et al., 2016; Parsons et al., 2010; Stark et al., 2019; Swain et al., 2014).

Focusing on infant cry perception, a meta-analysis confirmed the relevance of the networks in the parental brain by including 14 fMRI studies examining brain activity when listening to infant cry versus a matched control sound (Witteman et al., 2019). On the one hand, the meta-analysis confirmed the involvement of the auditory system, the thalamo-cingulate circuit, the dorsal anterior insula, the pre-supplementary motor area, dorsomedial prefrontal cortex and the inferior frontal gyrus (Witteman et al., 2019). On the other hand, the meta-analysis did not confirm the otherwise well-established involvement of the reward pathway in infant cry perception. The authors suggested that their result could have been due to sample size, the high-level control sound, or the use of unfamiliar and not own infant cry stimuli (Witteman et al., 2019). Furthermore, the study by Wittemann et al. (2019) only included fMRI studies, but left out other studies including Swain et al. (2014) and Young, Parsons, Stein, et al. (2017) that used electrophysiological methods to capture the temporal dynamics of brain activity in much smaller time-windows than fMRI, highlighting their potential to reveal the brain mechanisms underlying the parent-infant relationship. In addition to EEG, MEG represents an even more powerful tool to record brain activity due to its excellent temporal resolution and good spatial accuracy. Indeed, a study by Young and colleagues used MEG to measure brain activity when listening to infant versus adult cry in a group of non-parents (Young et al., 2016). This study found early differences in neural responses to infant versus adult cry in auditory, emotional and motor regions, including an early orbitofrontal cortex (OFC) response at around 130 and 195 ms. The OFC has been found to be an important hub for emotional, motivation and pleasure processing (Kringelbach, 2005; Kringelbach et al., 2016; Parsons, Young, Stein, et al., 2017) and this includes a fast response to infant faces at around 130ms (Kringelbach et al., 2008; Parsons, Young, Joensson, et al., 2014; Young et al., 2016). However, Young and colleagues primarily provided information on the brain mechanisms underlying infant cry processing in non-parents. Given that *anatomical* brain changes have been found in a longitudinal study of first-time mothers (Hoekzema et al., 2017), our study therefore focused on investigating the accompanying *functional* changes in brain responses to infant cry in a population composed of recent first-time mothers exhibiting different levels of bonding with their infants. To our knowledge, this is the first study to examine neural responses to infant cries in first-time mothers using MEG.

Equally, the experience of becoming a parent is often accompanied by a period of sleep deprivation linked to the arrival of the new family member. The sleep interruption that occurs during the first year of parenthood can be distressing, and studies have found neurobehavioral performance to decline during the postpartum period (Insana et al., 2013). In general, sleep deprivation has been found to affect both attention, memory, emotions, reward circuits, decision-making and resting-state networks by altering brain activity (Chee & Zhou, 2019; Krause et al., 2017; Wang et al., 2022), and lately two studies also found a link between lack of sleep and negative parenting (Chary et al., 2020; Leerkes et al., 2021). However, whether and how this impacts the parental brain is yet to be examined. Thus, in our study, we also assessed whether sleep deprivation has an impact on the brain responses to infant cry in new first-time mothers.

## Methods

### Data and code availability

On reasonable request, the codes and the anonymized neuroimaging data will be made available.

### Participants

Forty-three first-time mothers with infants ranging in age from 1.80 to 11.90 months (mean: 7.36 ± 2.03) were included. The participants varied in age from 23 to 42 years old (mean: 29.42 ± 3.65). Four participants were left-handed (9.1%), and four participants had undergone fertility treatment. All participants were recruited through social media websites like Facebook, local and national news outlets, and public bulletin boards. All participants were assessed for inclusion and exclusion criteria and had to sign the informed consent form before inclusion. Healthy first-time mothers over the age of 18 years were eligible to participate. A score of 11 or higher on the Edinburgh Postnatal Depression Scale (EPDS), comorbid psychiatric disorder, use of sleep medication, caffeine intake on the day of data collection, a history of sleep disorders, more than two nightshifts per week, and an uncorrected visual impairment were all used as exclusion criteria. The trial ran from August 2018 to February 2020 at Aarhus University Hospital, before being interrupted and prematurely ended by the COVID-19 pandemic.

The study was approved by the Ethics Committee of the Central Denmark Region (De Videnskabsetiske Komitéer for Region Midtjylland, approval number: 1-10-72-221-15). The Declaration of Helsinki - Ethical Principles for Medical Research was followed in all experimental procedures.

### Experimental design and stimuli

This is a cross-sectional neuroimaging study examining the relationship between brain activity when listening to infant cries compared to adult cries, total-night-time sleep, and postpartum bonding with the infant.

The experimental design is identical to that used in Young et al. (2016). Participants monitored and reported changes in the pitch of a control sound (500 ms of a 400 or 500 Hz tone) between trials in an implicit task, while infant vocalizations (laughter, cry, and neutral babble) and control sounds (adult cry, animal cry, and matched artificial infant laughter and cry) were delivered bi-aurally for 1500ms (see **Figure 1**). The implicit task was omitted from the analysis because its function was to keep the participants engaged in the experiment. The sounds were taken from the validated ‘OxVoc’ Sounds Database (Young, Parsons, LeBeau, et al., 2017). The ‘OxVoc’ Sounds Database is based on video recordings of infants interacting with their parents at home (Young et al., 2012). All infants were healthy, full-term, and between 6 and 8 months of age at the time of recording (M = 6.7 months, SD = 0.9). Adult cry stimuli were taken from video diary blogs, recorded by females at the age of 18–30 years (Parsons, Young, Craske, et al., 2014).

**Figure 1.**
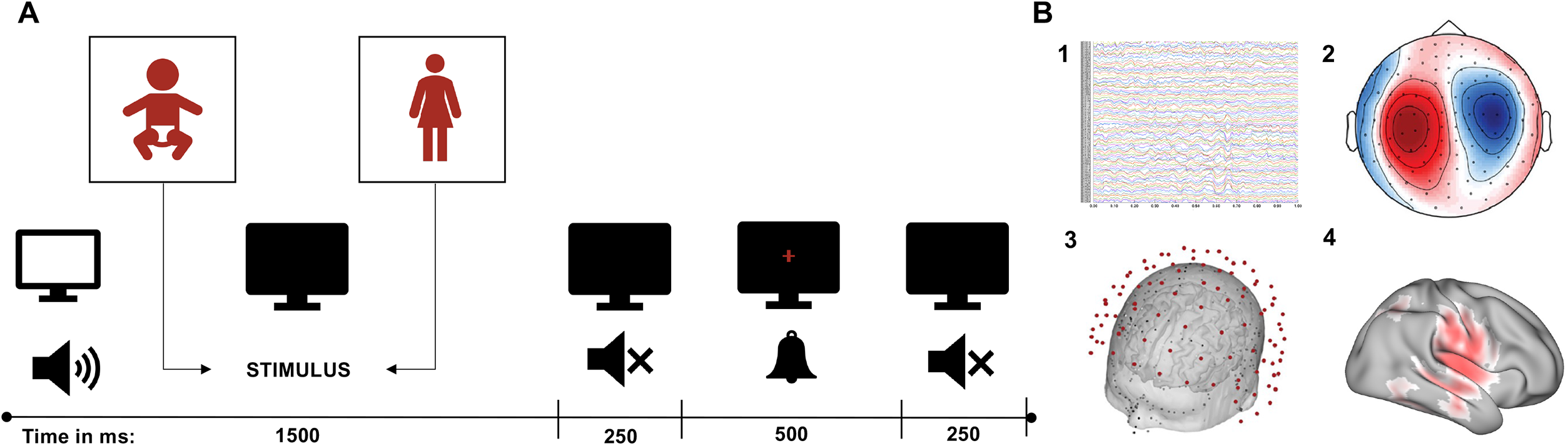
Overview of experimental design. **A)** The panel illustrates one trial from the paradigm used in the MEG. The total trial length is 2500 ms with an auditory stimulus of either an infant cry or an adult cry for 1500 ms. The stimulus was followed by 250 ms of silence, 500 ms of the implicit task sound (participants were asked to press a button when hearing a target sound (400 Hz)), and 250 ms of silence. Each category had 120 trials. **B)** The figure illustrates the MEG methods used in the further analysis. 1) Raw MEG signals were recorded. 2) Raw data was maxfiltered, transformed into SPM format and independent component analysis (ICA) were performed with manual artefact removal. 3) Data was beamformed and source reconstructed. 4) Finally, we contrasted infant cry versus adult cry, and correlated the postpartum bonding scores with brain activity. See Methods section for further details.

Each type of stimuli was repeated eight times, for a total of 120 trials per category. Presentation® Neurobehavioural Systems software was used to present all stimuli. MEG was utilised to measure brain activity during the presentation of the stimuli.

#### Questionnaires

Postpartum bonding was assessed using the Postpartum Bonding Questionnaire (Brockington et al., 2001). The PBQ is composed of 25 questions that are grouped into four categories: 1) Impaired bonding, 2) Rejection and pathological anger, 3) Infant-focused anxiety, and 4) Incipient abuse. We only focused on the domain of Impaired bonding (factor 1) in this study because this scale was the only one with a reasonable variance that allowed us to properly compute correlations in a later stage of our analysis pipeline. Factor 1 has 12 items with values ranging from 0 to 60, with higher scores indicating more impaired bonding. It has been proposed that a cut-off score of 12 indicates a high likelihood of impaired bonding (Brockington et al., 2006). Due to a late response on an ethical modification, the PBQ was incorporated after the trial began, resulting in a lesser quantity of PBQ-datasets (n = 29).

To avoid confounds from postnatal depression, we screened potential participants with the Edinburgh Postnatal Depression Scale (EPDS). This was used to evaluate the presence of postnatal depression symptoms using a Danish version (Smith-Nielsen et al., 2018). The EPDS is a 10-item questionnaire with a scale of 0 to 30, with higher scores indicating more depressive symptoms. When utilised in a Danish population, Smith-Nielsen et al. (2018) indicated a cut-off score of 11 or higher to determine whether there is a high risk of depression (Smith-Nielsen et al., 2018). Only participants with a score of less than 11 were included.

#### Sleep measurements

For sleep measurements, participants were given an actigraphy, GENEActiv original (Activinsights, UK), and a sleep log after being accepted into the study. An actigraphy is a movement-detecting accelerometer that is used to evaluate various sleep metrics, such as total night-time sleep. In addition, participants were required to complete a seven-day sleep journal. Actigraphy and sleep log measurements were recorded seven days before and after the MEG recording.

The GGIR package version 2.4-0 in R version 3.6.1 was used to analyse the actigraphy data (Migueles et al., 2019; van Hees et al., 2015). For details on the settings, see the **Supplementarty Table 3**. For the two weeks of measurements, the total amount of night-time sleep was averaged. For those who only had one week of actigraphy measurements, the average of that week was used (three participants in total).

### Neuroimaging data acquisition

Both the MRI and MEG data were collected in two separate sessions. An Elekta Neuromag TRIUX system (Elekta Neuromag, Helsinki, Finland) with 306 channels was used to collect the MEG data. The device is housed in a magnetically shielded room at Aarhus University Hospital in Denmark. The data was collected at a sampling rate of 1000 Hz with an analogue filtering of 0.1–330 Hz. Prior to the start of the experiment, the participants were asked to complete an auditory oddball test within the machine to ensure that the sound volume was 50 decibels above their minimum hearing threshold. We also used a 3D digitizer (Polhemus Fastrak, Colchester, VT, USA) to record the participant’s head shape and the position of four headcoils in relation to three anatomical landmarks (nasion, and left and right preauricular locations). We used this information to co-register the MEG data with the anatomical structure gathered by the MRI scanner during the study. Using a continuous head position identification system, the location of the headcoils was tracked during the recording (cHPI). While seated in the MEG scanner, we were able to trace the exact location of the participant’s head at each time point. We used this data to do an accurate movement correction at a later step of data analysis.

In the MRI, structural T1 data was collected. The following were the scan’s acquisition parameters: voxel size = 1.0 × 1.0 × 1.0 mm (or 1.0 mm3); reconstructed matrix size 256×256; echo time (TE) of 2.96 ms and repetition time (TR) of 5000 ms; and bandwidth of 240 Hz/Px. Using the Polhemus head shape data and the three fiducial points measured during the MEG session, each individual T1-weighted MRI scan was then co-registered to the standard Montreal Neurological Institute (MNI) brain template through an affine transformation and then referenced to the MEG sensors space.

### Data pre-processing

We used MaxFilter (Taulu & Simola, 2006) to pre-process the raw MEG sensor data (204 planar gradiometers and 102 magnetometers) for signal space separation to reduce interference from beyond the scalp. Maxfilter also downsampled the signal from 1000 Hz to 333 Hz and accounted for head movement.

The data was then transformed into the Statistical Parametric Mapping (SPM) format and processed in Matlab (MathWorks, Natick, Massachusetts, USA) using the Oxford Centre for Human Brain Activity Software Library (OSL) (M. Woolrich et al., 2011) and in-house-built functions. OSL is a open source MATLAB toolbox that combines custom functions with tools from the FMRIB Software Library (FSL) (M. W. Woolrich et al., 2009), SPM (Friston et al., 2007), and Fieldtrip (Oostenveld et al., 2011). The data was then filtered using a high-pass filter (0.1 Hz threshold). This was done in order to filter out frequencies that were too low to originate from the brain. We also used a 48-52 Hz notch filter to eliminate any potential electric current interference. We removed a few large artefacts from the data after visual inspection. An independent component analysis (ICA) was used to deconstruct the original signal into independent components in order to remove the interference of eyeblinks and heartbeat artefacts from the brain data. The components that detected eyeblink and heartbeat activities were then identified and removed, with the signal being rebuilt from the remaining components (Mantini et al., 2011). Finally, the data was epoched in 120 trials (per condition) of 1.850 milliseconds each (100ms of pre-stimulus time).

### Source reconstruction

#### Beamforming

Using OSL, we reconstructed the brain activity acquired on the scalp via MEG channels in source space (M. Woolrich et al., 2011). This was done using a forward model with local spheres and a beamformer approach as an inverse method (Hillebrand & Barnes, 2005) (see **Figure 1B**). In this beamforming implementation, we used both magnetometers and planar gradiometers, and a brain template with 8-mm resolution (resulting in 3559 brain sources or voxels). The spheres model represented the MNI-co-registered anatomy as a simpler geometric model, allowing each sensor to be fitted with its own sphere (Hillebrand & Barnes, 2005). Finally, the beamforming used a diverse set of weights sequentially applied to the source locations to isolate the contribution of each source (brain voxel) to the activity recorded by the MEG channels for each time-point in order to isolate the contribution of each source (brain voxel) to the activity recorded by the MEG channels for each time-point (Bonetti et al., 2020; Brookes et al., 2007; Hillebrand & Barnes, 2005).

#### Brain activity underlying listening to infant versus adult cry

After reconstructing the neural sources of the recorded brain data, we contrasted the brain activity underlying processing of infant versus adult cry. First, we sub-averaged the reconstructed brain signal in ten subsequent time-windows lasting 100ms each, from 1 to 1000ms from the onset of the stimuli (i.e., time-window one: 1 - 100ms; tw two: 101 – 200ms; tw three: 201-300ms, etc.). Second, we computed one paired-sample t-test for each reconstructed brain voxel. We corrected for multiple comparisons by using cluster-based Monte-Carlo simulations (MCS). Specifically, we computed the sizes of the clusters of significant neighbouring brain voxels (*p* < .005 from the paired-sample t-tests, corresponding to a t-value = 3. This cluster-forming threshold was obtained by dividing the standard **α** level = .05 by 10, which corresponds to the number of time-windows used in our study). Then we computed 1000 permutations of the original data. For each permutation, we computed the sizes of the clusters of significant neighbouring permuted brain voxels. By doing this 1000 times, we built a reference distribution of the biggest cluster sizes occurred in the permutated data. Then, we considered significant the original clusters that had a size bigger than the 99.9% of cluster sizes forming the reference distribution. Additional details on the MCS algorithm can be found in (Bonetti, Brattico, Bruzzone, et al., 2021; Bonetti, Brattico, Carlomagno, et al., 2021; Bonetti et al., 2020; Bruzzone et al., 2021).

#### Relationship between brain activity underlying listening to infant cry and bonding score

Next, we conducted a further analysis to assess whether the level of impaired bonding between the mother and her infant had an influence on the brain activity underlying processing of infant cry sounds. To achieve this aim, we computed Pearson’s correlations between the Postpartum Bonding Questionnaire (Brockington et al., 2001, 2006) score for each participant and the neural activity for each brain-voxel and time-window, followed by the same MCS described above (since this was a correlation analysis, our cluster-forming threshold was *r* = ± 0.4).

#### Relationship between brain activity underlying listening to infant cry and sleep disruption

To test for a correlation between neural activity and amount of objective average night-time sleep assessed through actigraphy measurements, we computed Pearson’s correlations between night-time sleep and neural activity. As previously done, we computed this analysis for each brain voxel and time-window. Finally, we corrected for multiple comparisons employing the same MCS as described above.

### Time-frequency analysis

In order to confirm the previous findings of a fast response in the OFC infant stimuli (Kringelbach et al., 2008; Parsons, Young, Stein, et al., 2017b; Young, Parsons, Stein, et al., 2017), we computed time-frequency analysis on the evoked responses of the OFC. Similar to the previous studies, we used complex Morlet wavelet transform (Daubechies & Heil, 1992), (from 8 to 40 Hz with 1-Hz intervals, as carried out by Kringelbach et al. (2008)) and carried out two independent analyses for both left and right OFC. We then contrasted the power spectra of infant versus adult cry. We did that for each frequency, hemisphere, and time-point within the range: 0 –400 ms, similar to previous studies (Kringelbach et al., 2008; Parsons, Young, Stein, et al., 2017b; Young, Parsons, Stein, et al., 2017). The emerging *p*-values were binarized according to the threshold a = .05 and then submitted to a two-dimensional MCS. Here, we computed the clusters size of continuous significant values in time and frequency. Then, we made 10000 permutations of the binarized *p*-values. For each permutation, we extracted the size of the maximum cluster and built a reference distribution with one value for each permutation. Finally, the original clusters were considered significant if they had a size bigger than the 99.99% of the maximum cluster sizes of the permuted data.

## Results

### Experimental design and data analysis overview

We used MEG to investigate the later stages of brain processing in new first-time mothers hearing vocalisations, specifically comparing their processing of infant and adult cry vocalisations. We were also interested in investigating if this brain activity was correlated with levels of non-clinically significant sleep deprivation (actigraphy measurements and sleep logs) and impaired bonding (Postpartum Bonding Questionnaire). The MEG data was pre-processed and analysed in Matlab (MathWorks, Natick, Massachusetts, USA) following the OSL pipeline. After source reconstruction, we contrasted infant versus adult cry on a whole-brain level within ten different non-overlapping time-windows. Finally, in two further analyses, we correlated the neural activity with total night-time sleep and the impaired bonding score on a whole-brain level within the ten time-windows.

### Late brain activity underlying listening to infant versus adult cry

After reconstruction of the brain sources of the MEG sensor level signal, we contrasted the neural activity underlying processing of infant versus adult cry, independently for each of the 3559 brain voxels and each of the ten time-windows considered in the study (see Methods for details).

The analysis, corrected for multiple comparisons by employing cluster based MCS, are reported in **Table 1, Supplementary Tables 1 and 2**, and returned an array of significant clusters. Specifically, we detected significant clusters (*p* < .001, k cluster size = 219 and k = 293, respectively) for the last two time-windows (801-900ms and 901-1000ms from the onset of the stimulus). As depicted in **Figure 2**, results indicated that processing infant versus adult cry sounds elicited a stronger brain signal, especially in brain regions located in the right hemisphere such as primary auditory cortex, superior temporal gyrus, hippocampal areas, insula, post-central gyrus, supramarginal gyrus and posterior cingulate gyrus.

**Table 1:**
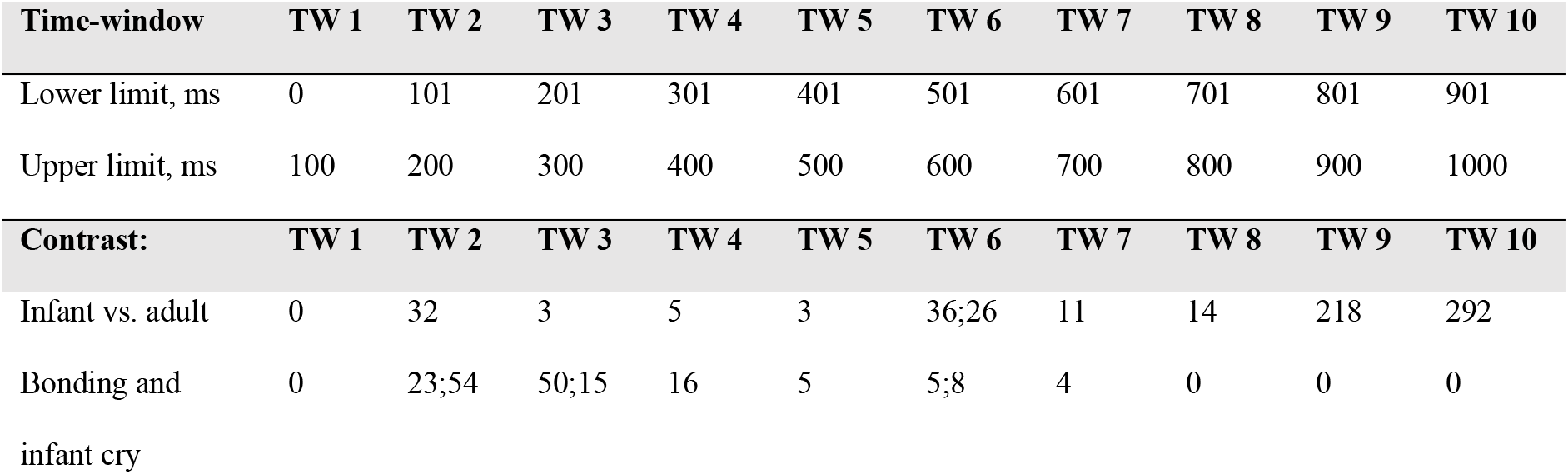
Cluster size for the different time-window(s)

**Figure 2.**
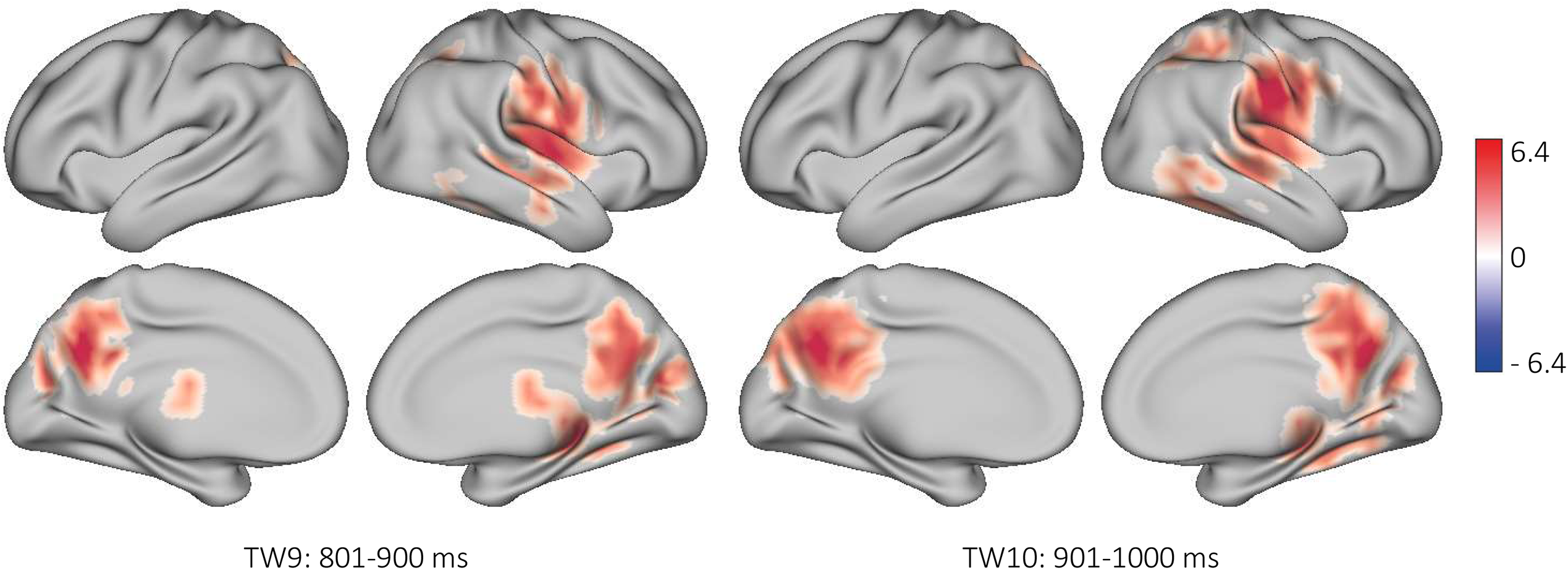
Late brain differences in processing infant versus adult cry. This figure shows the clusters of significantly higher activity when listening to infant cries compared to adult cries after 800 ms. These are the clusters seen in the late time-windows, and show statistically significant increased activity especially in the right hemisphere, in the anatomical areas such as primary auditory cortex, superior temporal gyrus, hippocampal areas, insula, precuneus supramarginal gyrus, post-central gyrus and posterior cingulate gyrus.

### Relationship between brain activity underlying listening to infant cry and bonding score

The PBQ score ranged from 1 to 18 with only 2 participants having a score above 12 indicating a high risk of impaired bonding. The mean PBQ score was 6.69 ± 3.82 (n = 29). We computed an additional analysis to assess whether the impaired bonding score from the PBQ affected the brain processing of infant cries. We computed Pearson’s correlation between bonding score and the brain activity and corrected for multiple comparisons using the same MCS as described above. As reported in **Table 1**, results showed significant clusters especially in the time-windows two and three (i.e., 101-200ms and 201-300ms; *p* < .001, tw two: k = 24, k = 55, tw three: k = 51, k = 16). These clusters, shown in **Figure 3** show that a lower bonding with the infant (meaning higher impaired bonding score) was associated with a reduced activity in brain regions such as primary auditory cortex, middle and superior temporal gyrus, medial OFC, hippocampal areas, supramarginal gyrus and inferior frontal gyrus.

**Figure 3.**
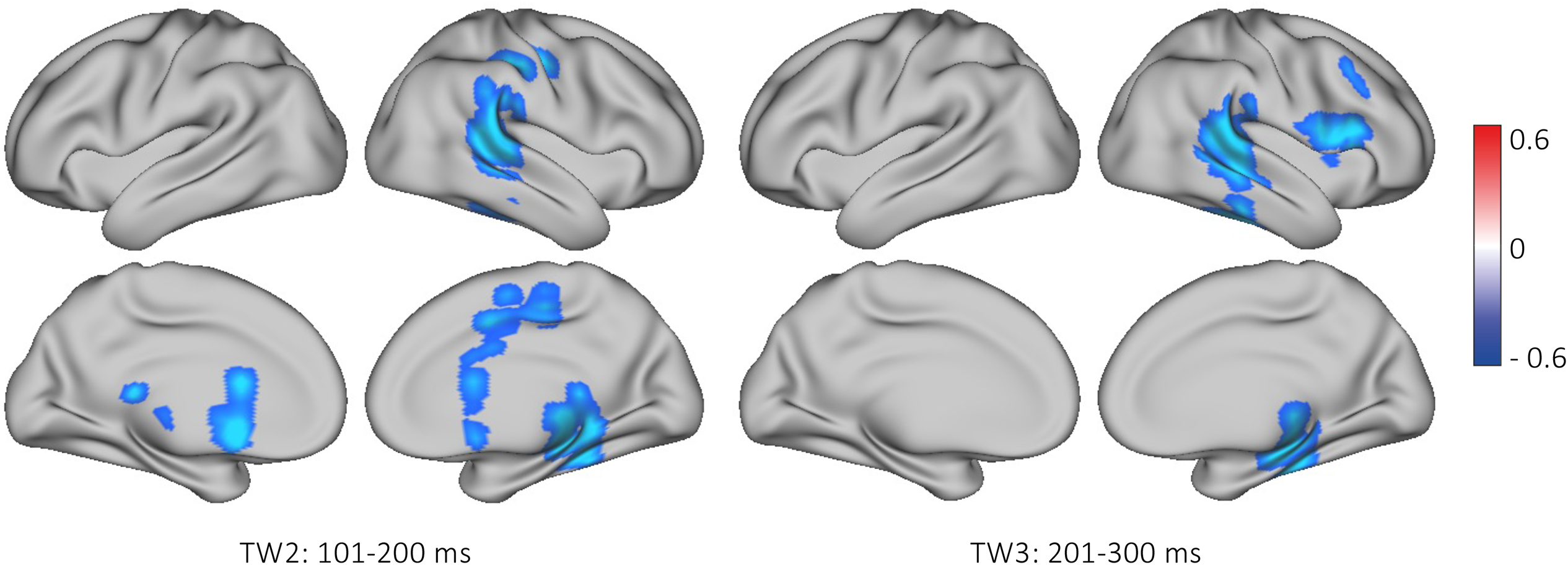
Weaker bonding scores correlate with early activity in brain regions. The figure shows the clusters with significantly lower activity when correlating neural activity to infant cries and weaker bonding scores. The higher a score, the more severe is the impaired bonding. These are the clusters seen in time-windows (TW) 2 and 3. The correlation of neural activity and impaired bonding showed statistically significant reductions in neural activity in the anatomical areas including the primary auditory cortex, superior and middle temporal gyrus, medial OFC, hippocampal areas, supramarginal gyrus and inferior frontal gyrus, most strongly in the right hemisphere. This could suggest an early obstacle to successful bonding.

### Correlation between brain activity underlying listening to infant cry and sleep disruption

On average, the mothers slept 7.15 ± 0.76 hours of night-time sleep. There was no statistically significant correlation between the amount of average night-time sleep and the MEG signals.

### Fast response in OFC determined by time-frequency analysis

Finally, we ascertained that the MEG data also included a fast response to infant cries as reported earlier (Young et al. (2016)). We computed time-frequency analysis on the evoked responses of the OFC and contrasted the power spectra for infant versus adult cry. As specified in the Methods, analyses were conducted separately for left and right OFC and multiple comparisons were corrected by using cluster-based MCS. Results returned four significant clusters, all localized in the left hemisphere. As can be seen in **Figure 4**, this analysis confirmed the earlier findings with stronger power for infant versus adult cries for 8 Hz and in the time range: 120-250 milliseconds. Interestingly, later on there were three clusters showing the opposite tendency, namely a stronger power for adult versus infant cries. They were found for the following frequency and time-ranges: 17-19 Hz and 300-380 milliseconds (i); 29-34 Hz and 210-250 milliseconds (ii); 16 Hz and 190-260 milliseconds (iii). The complete contrasts for the power spectra of infant versus adult cries are shown in **Figure 4**.

**Figure 4.**
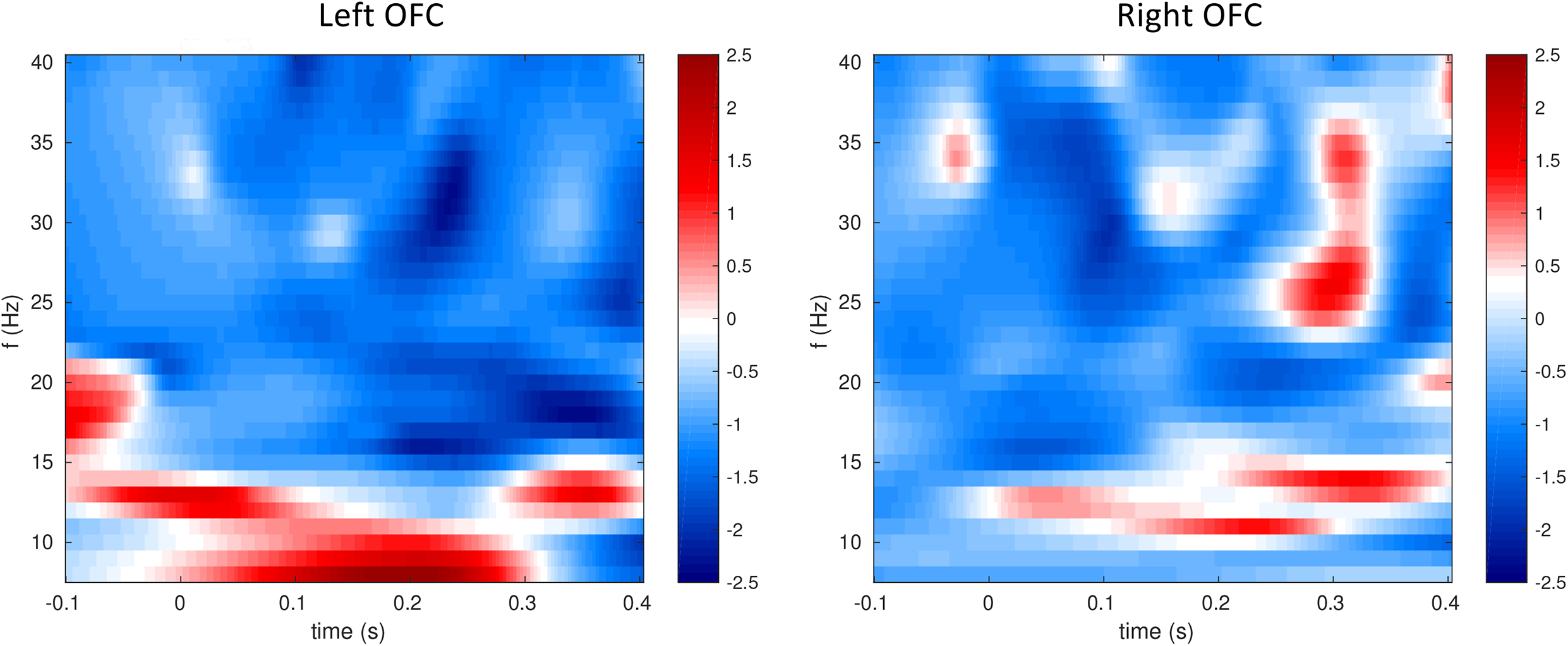
Early activity in the orbitofrontal cortex (OFC) to infant cries. The figure shows the contrast of the power spectra for infant versus adult cry in the left OFC (left panel) with significant differences from around 120 to 250 ms after stimulus onset around 8-10 Hz for infant versus adult cry. This confirms the findings of earlier studies (Kringelbach et al., 2008; Parsons, Young, Stein, et al., 2017b; Young, Parsons, Stein, et al., 2017).

## Discussion

In this study we investigated the temporal evolution of brain activity in new first-time mothers processing infant and adult cries. We also investigated if naturally occurring differences in 1) sleep deprivation and 2) individual differences in bonding with the infant in the first-time mothers correlated with their brain activity.

Using a time-windowed approach, we found significant differences in the processing of infant versus adult cry sounds in the late stages of brain activity after around 800ms after the onset of each vocalisation lasting 1500ms. The changes were found especially in primarily right hemisphere regions including primary auditory cortex, precuneus, supramarginal gyrus, superior temporal gyrus, hippocampal areas, insula, post-central gyrus and posterior cingulate gyrus. Furthermore, correlating of the windowed brain activity with weaker bonding scores revealed statistically significant reductions in neural activity regions including in the primary auditory cortex, medial OFC, hippocampal areas, supramarginal and inferior frontal gyrus. In contrast, we did not find a statistically significant correlation between the windowed brain activity and sleep deprivation scores.

In contrast to our earlier studies, which were designed to reveal the very fast brain processing of infant stimuli (Kringelbach et al., 2008; Parsons, Young, Joensson, et al., 2014; Young et al., 2016), here we took a different, slower approach to investigate the later stage of the temporal evolution of brain processing. We therefore used a time-windowed approach to MEG analysis which is optimised to find these later changes. Our finding of later significant changes around 800ms after onset of vocalisations revealed increased neural activity in the mentalisation and empathy networks (precuneus, superior temporal gyrus, posterior cingulate gyrus, supramarginal gyrus), along with activity in emotional regulation and evaluation regions (superior temporal gyrus, insula) (Bjertrup et al., 2019; Feldman, 2015; Swain et al., 2014; Witteman et al., 2019). These changes in functional activity fits well with the previously identified longitudinal structural brain changes (Hoekzema et al., 2017) and shed new light on learning-related changes in caregiving.

The correlation between the brain activity related to infant cries and postpartum bonding score showed decreased activity after 100-200ms in primary auditory cortex, medial OFC, hippocampal areas, supramarginal gyrus, and inferior frontal gyrus. This is of considerable interest given the known role of the medial OFC in caregiving (Kringelbach et al., 2016). This could indicate that the degree of bonding with the infant is directly correlated with activity in this key region of the parental brain.

Equally, the inferior frontal gyrus and supramarginal gyrus are also part of the mentalisation and empathy networks important for the parental brain. Evidence has linked inferior frontal gyrus as part of a network that has been shown to be especially important for understanding another person’s intentions (Bjertrup et al., 2019; Feldman, 2015; Schurz et al., 2014; Swain et al., 2014; Witteman et al., 2019), and the right supramarginal gyrus has been found important in the correct assessment of other people’s emotions despite one’s own emotional state (Silani et al., 2013).

In a study by Hoekzema et al. (2017), they followed the longitudinal anatomical changes from before pregnancy to two years postpartum (Hoekzema et al., 2017). The results showed grey matter volume decreases in a pattern resembling the theory-of mind network. The reduction in grey matter was interpreted as a maturation process, and they found increased activity in these areas when mothers were watching pictures of their infants during fMRI (Hoekzema et al., 2017). To understand how significant the changes of the maternal brain were compared to a non-parent female brain, Hoekzema et al. (2017) used a multikernel learning approach to find the areas of highest predictive power in deciding whether the brain in question belonged to a mother or a non-mother. Here they found the right middle temporal gyrus, inferior frontal gyrus and posterior cingulate cortex to be of highest predictive power (Hoekzema et al., 2017). Hence, our results suggest that the decreased functional activity in the inferior frontal gyrus and right middle temporal gyrus correlating with the higher impaired bonding score might represent a problem with the maturation of the ‘parental brain’ network.

Furthermore, the decreased activity in the primary auditory cortex might reflect a decreased sensitivity to the infant cry. Maternal sensitivity has been found to be affected by the mother’s own childhood experience, i.e., if you had a warm secure maternal attachment in childhood, you are more likely to become a warm sensitive mother yourself providing a secure attachment style to your child (Belsky et al., 2005). A neuroimaging study by Kim and colleagues (2010) found that new mothers reporting maternal warmth in their own experienced childhood had increased activity in emotional regulatory areas such as the middle and superior temporal gyri, while others reporting less maternal warmth in their own childhood experience showed increased left hippocampal activations when listening to infant cries (Kim et al., 2010). The study by Kim et al. (2010) used a retrospective self-measurement of paternal bonding experienced in their childhood, while the present study examined how brain activity correlated with their own perceived bonding with the infant, which might explain the opposite effects on hippocampal neural activity. Moreover, in our study, only two mothers had a Postpartum Bonding score above the cut-off score for high risk of impaired bonding. Thus, the neural activity in a population of mothers with even higher scores might vary from the results reported in our study. Nevertheless, the observed decreases in neural activity correlating with higher scores of impaired bonding in a population of healthy mothers might suggest a neural signature of risk of impaired bonding. Furthermore, we confirmed the fast OFC response after around 130ms previously seen in young non-parents (Kringelbach et al., 2008; Parsons, Young, Stein, et al., 2017b; Young, Parsons, Stein, et al., 2017). Hence, we suggest that that a natural learning process takes place when becoming a parent for the first time, developing this innate ‘parental instinct’ into higher cognitive processing of infant stimuli. However, a longitudinal study following couples before and after pregnancy using MEG investigating both early and late perceptual processing of infant cues is needed to fully conclude on this.

Finally, we examined whether neural activity and night-time sleep were correlated, since sleep deprivation (both acute and chronic) has been found to impact brain activity regarding attention, memory, emotions, reward circuits, decision-making, and resting-state networks (Chee & Zhou, 2019; Krause et al., 2017; Wang et al., 2022). Thus, sleep deprivation affects several important networks for eliciting and conducting appropriate caregiving behaviour. Nonetheless, in our sample of non-significant sleep deprived mothers, we did not find a correlation between level of night-time sleep and neural activity involved in the perception of infant cries. Perhaps infant stimuli are ‘super salient’ due to the evolutionary importance of infant survival and thus the brain mechanisms underlying their processing may not be dramatically impaired by moderate levels of sleep deprivation. This suggestion is backed up by a recent study by Hoegholt et al. (in submission). In this study, we examined the effect of night-time sleep on the self-prioritization effect in new first-time mothers (Hoegholt et al. in submission). The self-prioritization effect has been established over several cognitive domains (attention, perception, memory and decision making), and refers to the bias of being more responsive to stimuli referring to self, as when hearing one’s own name in a crowd of loud people while engaged in another conversation (Moray, 1959; Sui et al., 2012; Sui & Humphreys, 2015; Sui & Rotshtein, 2019; Yin et al., 2019). Hoegholt et al. divided the new mothers into two groups depending on their average night-time sleep, above or below 7 hours, and found that the ones sleeping above 7 hours prioritized self and infant stimuli equally, while those that slept less than 7 hours prioritized infant above all other stimuli (Hoegholt et al. in submission).

There are some potential limitations to the current study. We failed to observe the previously finding of changes in amygdala activity where the duration of motherhood has been found to increase activity in this region (Parsons, Young, Petersen, et al., 2017). However, this study used fMRI and not MEG (Parsons, Young, Petersen, et al., 2017), as do most of the studies the parental brain models are based on (Feldman, 2015; Swain et al., 2014; Witteman et al., 2019). While MEG has great temporal resolution in cortical regions, it is less sensitive to activity in subcortical regions. Another potential limitation to this study was the sudden disruption caused by the COVID-19 pandemic, as more participants might have allowed for a more sensitive analysis of the Postpartum Bonding Questionnaire scores and sleep levels with brain activity. We suggest future studies to include both fMRI and MEG/EEG methods to advance our understanding of the temporal dynamics of the parental brain. Moreover, behavioural measures of the parent-infant relationship are needed to establish whether impaired bonding and hence the risk of neglect can be predicted by a neural signature.

## Conclusion

We applied the experimental paradigm by Young et al (2016) on a new population of first-time mothers within the first 12 months of motherhood, measuring both night-time sleep and self-reported levels of parent-infant bonding.

In accordance with the proposed models of the parental brain, we found increased neural activity for infant cry in the mentalisation and empathy networks (e.g. precuneus, superior temporal gyrus, posterior cingulate gyrus, supramarginal gyrus), along with activation in some emotional regulation and evaluation regions (superior temporal gyrus, insula). Furthermore, we demonstrated that reduced early brain responses to infant crying could be a biomarker of a potential mother-infant bonding problem. Finally, the results could suggest that infant crying is of such high importance that sleep deprivation has no effect on the perception of this crucial stimulus. However, more research is needed to draw firm conclusions from these findings.

## Acknowledgements

Special thanks are given to all the student researchers that helped out with the data collection.

This work was funded by Center for Music in the Brain (MIB) which is funded by the Danish National Research Foundation (project number DNRF117); and a European Research Council Consolidator Grant CAREGIVING (grant number: 615539) to MLK.

LB is supported by Carlsberg Foundation (CF20-0239), Center for Music in the Brain, Linacre College of the University of Oxford, and Society for Education and Music Psychology (SEMPRE’s 50th Anniversary Awards Scheme).

MLK is supported by the ERC Consolidator Grant: CAREGIVING (n. 615539), Center for Music in the Brain, and Centre for Eudaimonia and Human Flourishing funded by the Pettit and Carlsberg Foundations.

Address of corresponding author: 7 Stoke Place, OX3 9BX Oxford, UK

## Author contributions

NFH, ABAS, HMF and MLK conceived the hypotheses and designed the study. NFH, ABAS, HMF and CA were responsible for the data-collection. LB, NFH and CA performed the pre-processing and statistical analysis of the MEG data, while NFH, MH and CA pre-processed and analysed the sleep data. MLK, PV, ABAS, LB and HMF provided essential help to interpret and frame the results within the neuroscientific literature. NFH wrote the first draft of the manuscript and, together with LB, prepared the figures. All the authors contributed to and approved the final version of the manuscript.

