## Supplementary material for "A potential marker for problematic mother-infant bonding revealed by magnetoencephalography study of first-time mothers listening to infant cries"

### SUPPLEMENTARY TABLES

***Table ST1 – Detailed information on significant clusters for MEG sensor data comparing infant versus adult cries***

*MCS revealed significant clusters of MEG sensors that contrasted infant versus adult cries. Those clusters are depicted in the table in terms of significant channels and time-windows.*

***Table ST2 – Detailed information on significant clusters for MEG sensor data correlating activity with impaired bonding***

*The Pearsons correlation showed significant clusters of decreased activity in several clusters, when correlating the impaired bonding score with neural activity. Those clusters are depicted in the table in terms of significant channels and time-windows.*

***Table ST3 – Actigraphy setting***

*An overview of the settings selected for the analysis of the actigraphy data.*

### Table ST1

**Time window 2, cluster, infant vs. adult cry**

| Brain Region | Hemisphere | t-value | MNI Coordinates *x* | *y* | *z* |
|---|---|---|---|---|---|
| Postcentral | R | 4.4 | 50 | -14 | 48 |
| Parietal Inf | R | 3.98 | 58 | -22 | 48 |
| Postcentral | R | 3.77 | 58 | -22 | 32 |
| Parietal Inf | R | 3.77 | 58 | -22 | 40 |
| SupraMarginal | R | 3.61 | 66 | -22 | 32 |
| Temporal Sup | R | 3.59 | 50 | -30 | 16 |
| Temporal Sup | R | 3.51 | 58 | -30 | 16 |
| Rolandic Oper | R | 3.5 | 42 | -30 | 16 |
| SupraMarginal | R | 3.49 | 58 | -30 | 32 |
| Postcentral | R | 3.43 | 66 | -14 | 32 |
| Temporal Sup | R | 3.36 | 50 | -30 | 8 |
| SupraMarginal | R | 3.35 | 50 | -30 | 24 |
| Precentral | R | 3.35 | 34 | -6 | 56 |
| SupraMarginal | R | 3.34 | 66 | -22 | 40 |
| Postcentral | R | 3.32 | 50 | -22 | 32 |
| Heschl | R | 3.27 | 42 | -30 | 8 |
|  | R | 3.25 | 34 | -30 | 16 |
| Parietal Inf | R | 3.25 | 58 | -30 | 40 |
|  | R | 3.23 | 26 | -6 | 48 |
| Postcentral | R | 3.22 | 42 | -22 | 40 |
| Frontal Sup | R | 3.21 | 26 | -6 | 56 |
| Postcentral | R | 3.19 | 50 | -22 | 40 |
| SupraMarginal | R | 3.18 | 66 | -30 | 32 |
|  | R | 3.18 | 42 | -30 | 24 |
| Temporal Sup | R | 3.17 | 58 | -30 | 8 |
|  | R | 3.14 | 50 | -22 | 24 |
| Precentral | R | 3.12 | 34 | -6 | 48 |
| Postcentral | R | 3.11 | 42 | -14 | 40 |
| Postcentral | R | 3.06 | 50 | -14 | 40 |
| Postcentral | R | 3.06 | 66 | -22 | 24 |
| Precentral | R | 3.06 | 42 | -14 | 48 |
| Postcentral | R | 3.04 | 58 | -14 | 32 |

**Time window 3, cluster, infant vs. adult cry**

| **Brain Regic** | **Hemispher** | **t-value** | **MNI Coordinates** | | |
|---|---|---|---|---|---|
| | | | ***x*** | ***y*** | ***z*** |
| | R | 3.21 | 26 | -38 | -32 |
| | R | 3.12 | 34 | -46 | -32 |
| | R | 3.08 | 26 | -38 | -40 |

**Time window 4, cluster, infant vs. adult cry**

| **Brain Regic** | **Hemispher** | **t-value** | **MNI Coordinates** | | |
|---|---|---|---|---|---|
| | | | ***x*** | ***y*** | ***z*** |
| | R | 3.24 | 26 | -62 | 0 |
| Fusiform | R | 3.2 | 34 | -62 | -8 |
| Fusiform | R | 3.11 | 34 | -54 | -8 |
| Lingual | R | 3.1 | 18 | -62 | 0 |
| | R | 3.06 | 42 | -54 | -8 |

**Time window 5, cluster, infant vs. adult cry**

| **Brain Region** | **Hemisphere** | **t-value** | **MNI Coordinates** | | |
|---|---|---|---|---|---|
| | | | ***x*** | ***y*** | ***z*** |
| SupraMargina | L | 3.34 | -54 | -30 | 32 |
| SupraMargina | L | 3.24 | -54 | -30 | 24 |
| | L | 3.06 | -46 | -30 | 32 |

**Time window 6, cluster 1, infant vs. adult cry**

| Brain Region | Hemisphere | t-value | MNI Coordinates | | |
|---|---|---|---|---|---|
| | | | *x* | *y* | *z* |
| | L | 3.8 | -14 | -62 | 40 |
| | R | 3.78 | 2 | -54 | 8 |
| Precuneus | L | 3.72 | -6 | -62 | 32 |
| | R | 3.65 | 2 | -62 | 8 |
| Cuneus | L | 3.63 | -6 | -86 | 40 |
| Precuneus | L | 3.63 | -6 | -62 | 40 |
| Precuneus | L | 3.57 | -6 | -54 | 32 |
| | L | 3.51 | -14 | -54 | 40 |
| | L | 3.44 | -14 | -46 | 24 |
| Cuneus | L | 3.42 | -14 | -62 | 32 |
| Precuneus | L | 3.38 | -6 | -54 | 40 |
| Parietal Sup | L | 3.36 | -14 | -62 | 48 |
| Precuneus | L | 3.36 | -6 | -62 | 48 |
| Precuneus | L | 3.31 | -6 | -70 | 32 |
| Precuneus | L | 3.3 | -6 | -62 | 24 |
| Precuneus | L | 3.3 | -6 | -78 | 56 |
| Calcarine | L | 3.29 | -6 | -62 | 16 |
| Precuneus | L | 3.28 | -14 | -70 | 40 |
| | L | 3.27 | -14 | -54 | 32 |
| Precuneus | L | 3.25 | -14 | -70 | 48 |
| Lingual | L | 3.24 | -6 | -62 | 8 |
| Precuneus | L | 3.24 | -6 | -70 | 40 |
| Parietal Sup | L | 3.2 | -14 | -78 | 56 |
| Precuneus | L | 3.2 | -6 | -54 | 24 |
| | R | 3.17 | 2 | -62 | 0 |
| Cingulum Mic | L | 3.17 | -6 | -46 | 32 |
| Precuneus | L | 3.16 | -6 | -54 | 48 |
| Precuneus | L | 3.16 | -6 | -70 | 48 |
| Precuneus | L | 3.15 | -6 | -78 | 48 |
| Precuneus | L | 3.14 | -6 | -70 | 56 |
| Precuneus | L | 3.13 | -6 | -62 | 56 |
| | L | 3.05 | -14 | -54 | 48 |
| | R | 3.05 | 2 | -54 | 0 |
| Precuneus | L | 3.03 | -6 | -46 | 40 |
| Precuneus | L | 3.01 | -6 | -70 | 64 |
| Lingual | L | 3.01 | -6 | -62 | 0 |

**Time window 6, cluster 2, infant vs. adult cry**

| Brain Region | Hemisphere | t-value | MNI Coordinates | | |
|---|---|---|---|---|---|
| | | | *x* | *y* | *z* |
| Heschl | R | 4.79 | 50 | -14 | 8 |
| Precentral | R | 4.55 | 50 | -6 | 24 |
| Insula | R | 4.37 | 42 | -14 | 8 |
| Rolandic Oper | R | 4.27 | 50 | -14 | 16 |
| Postcentral | R | 4.21 | 58 | -6 | 24 |
| Heschl | R | 4.16 | 58 | -14 | 8 |
| Insula | R | 4.11 | 42 | -14 | 16 |
| Postcentral | R | 4.04 | 58 | -14 | 16 |
| Rolandic Oper | R | 3.96 | 42 | -6 | 16 |
| Rolandic Oper | R | 3.89 | 50 | -6 | 16 |
| Rolandic Oper | R | 3.78 | 50 | -6 | 8 |
| | R | 3.71 | 42 | -6 | 24 |
| Temporal Sup | R | 3.71 | 66 | -14 | 8 |
| Precentral | R | 3.65 | 58 | -6 | 32 |
| Insula | R | 3.64 | 42 | -6 | 8 |
| Rolandic Oper | R | 3.63 | 58 | -6 | 8 |
| Postcentral | R | 3.6 | 66 | -14 | 16 |
| Postcentral | R | 3.53 | 58 | -6 | 16 |
| Rolandic Oper | R | 3.38 | 66 | -6 | 8 |
| Postcentral | R | 3.33 | 66 | -6 | 32 |
| Postcentral | R | 3.3 | 66 | -6 | 24 |
| Temporal Sup | R | 3.29 | 50 | -14 | 0 |
| | R | 3.25 | 34 | -14 | 16 |
| Putamen | R | 3.24 | 34 | -14 | 8 |
| Heschl | R | 3.14 | 42 | -22 | 8 |
| Precentral | R | 3.03 | 50 | -6 | 32 |

**Time window 7, cluster, infant vs. adult cry**

| **Brain Region** | **Hemisphere** | **t-value** | **MNI Coordinates** | | |
|---|---|---|---|---|---|
| | | | ***x*** | ***y*** | ***z*** |
| Precuneus | L | 4.17 | -6 | -54 | 40 |
| Precuneus | L | 3.99 | -6 | -62 | 32 |
| Precuneus | L | 3.75 | -6 | -62 | 24 |
| Precuneus | R | 3.56 | 2 | -62 | 48 |
| Occipital Sup | L | 3.39 | -22 | -62 | 32 |
| | L | 3.28 | -14 | -54 | 40 |
| Calcarine | R | 3.25 | 2 | -62 | 16 |
| Precuneus | R | 3.22 | 2 | -70 | 40 |
| | R | 3.16 | 2 | -54 | 16 |
| Precuneus | L | 3.06 | -14 | -62 | 24 |
| | L | 3.05 | -14 | -54 | 32 |

**Time window 8, cluster, infant vs. adult cry**

| Brain Region | Hemisphere | t-value | MNI Coordinates | | |
|---|---|---|---|---|---|
| | | | *x* | *y* | *z* |
| Precuneus | L | 4.48 | -6 | -62 | 32 |
| Precuneus | L | 4.46 | -6 | -62 | 24 |
| Precuneus | L | 4.15 | -14 | -62 | 24 |
| Calcarine | L | 4 | -6 | -62 | 16 |
| Precuneus | L | 3.83 | -14 | -54 | 24 |
| | L | 3.7 | -14 | -54 | 40 |
| Calcarine | R | 3.65 | 2 | -62 | 16 |
| Precuneus | L | 3.65 | -6 | -54 | 40 |
| | L | 3.6 | -22 | -54 | 32 |
| Precuneus | R | 3.58 | 2 | -70 | 32 |
| Precuneus | L | 3.49 | -6 | -54 | 16 |
| | L | 3.44 | -14 | -46 | 24 |
| Precuneus | R | 3.18 | 2 | -46 | 48 |
| | L | 3.15 | -22 | -54 | 40 |

**Time window 9, cluster, infant vs. adult cry**

| Brain Region | Hemisphere | t-value | MNI Coordinates | | |
|---|---|---|---|---|---|
| | | | *x* | *y* | *z* |
| | R | 5.88 | 34 | -30 | 0 |
| | R | 5.43 | 26 | -30 | 8 |
| | R | 5.15 | 34 | -30 | 8 |
| Cuneus | R | 5.03 | 2 | -70 | 24 |
| Precuneus | L | 5 | -6 | -62 | 32 |
| | R | 4.89 | 26 | -30 | 0 |
| Rolandic Oper | R | 4.87 | 50 | -6 | 16 |
| Insula | R | 4.8 | 42 | -14 | 16 |
| Insula | R | 4.76 | 42 | -14 | 8 |
| Thalamus | R | 4.73 | 18 | -30 | 8 |
| Rolandic Oper | R | 4.73 | 50 | -14 | 16 |
| Heschl | R | 4.72 | 42 | -22 | 8 |
| Postcentral | R | 4.69 | 58 | -6 | 16 |
| Hippocampus | R | 4.68 | 34 | -38 | 0 |
| Temporal Sup | R | 4.67 | 66 | -14 | 8 |
| Rolandic Oper | R | 4.67 | 58 | -6 | 8 |
| Hippocampus | R | 4.67 | 26 | -38 | 0 |
| Rolandic Oper | R | 4.64 | 50 | -6 | 8 |
| Precuneus | L | 4.62 | -6 | -54 | 32 |
| Precuneus | L | 4.57 | -6 | -62 | 40 |
| Rolandic Oper | R | 4.55 | 42 | -22 | 16 |
| Cingulum Mid | R | 4.54 | 2 | -54 | 32 |
| Precuneus | R | 4.53 | 2 | -62 | 24 |
| Insula | R | 4.5 | 42 | -14 | 0 |
| Heschl | R | 4.5 | 58 | -14 | 8 |
| | R | 4.48 | 34 | -22 | 0 |
| Rolandic Oper | R | 4.47 | 42 | -6 | 16 |
| Cuneus | R | 4.46 | 2 | -78 | 24 |
| Postcentral | R | 4.45 | 50 | -22 | 32 |
| | R | 4.42 | 42 | -30 | 0 |
| Precuneus | R | 4.42 | 2 | -62 | 32 |
| Postcentral | R | 4.39 | 58 | -14 | 32 |
| Precuneus | L | 4.37 | -6 | -70 | 32 |
| Hippocampus | R | 4.36 | 34 | -30 | -8 |
| Postcentral | R | 4.36 | 58 | -14 | 16 |
| Insula | R | 4.36 | 42 | -6 | 8 |
| | R | 4.35 | 42 | -38 | -8 |
| Calcarine | R | 4.35 | 2 | -62 | 16 |
| Temporal Sup | R | 4.33 | 50 | -22 | 8 |
| Thalamus | R | 4.32 | 26 | -22 | 0 |
| Rolandic Oper | R | 4.31 | 66 | -6 | 8 |
| Precuneus | R | 4.31 | 2 | -54 | 24 |
| Cuneus | L | 4.3 | -6 | -70 | 24 |
| | R | 4.29 | 42 | -30 | -8 |
| ParaHippocamp | R | 4.29 | 34 | -38 | -8 |
| Thalamus | R | 4.27 | 18 | -30 | 0 |
| Calcarine | R | 4.27 | 2 | -70 | 16 |
| | R | 4.25 | 34 | -22 | 8 |
| Precuneus | R | 4.23 | 2 | -54 | 40 |

| | | | | | |
|---|---|---|---|---|---|
| Postcentral | R | 4.19 | 66 | -6 | 16 |
| | R | 4.18 | 50 | -14 | 24 |
| | R | 4.13 | 26 | -22 | 8 |
| | R | 4.11 | 42 | -14 | 24 |
| Precuneus | L | 4.07 | -6 | -62 | 24 |
| Heschl | R | 4.07 | 42 | -30 | 8 |
| Rolandic Oper | R | 4.07 | 50 | -22 | 16 |
| Postcentral | R | 4.06 | 58 | -6 | 24 |
| Lingual | R | 4.06 | 18 | -38 | 0 |
| | R | 4.04 | 10 | -30 | 0 |
| Precuneus | L | 4.02 | -6 | -54 | 24 |
| | R | 4.02 | 42 | -22 | 0 |
| Putamen | R | 4.01 | 34 | -14 | 8 |
| | R | 4 | 2 | -54 | 16 |
| | R | 3.99 | 34 | -22 | 16 |
| Temporal Sup | R | 3.98 | 50 | -14 | 0 |
| | L | 3.98 | -14 | -62 | 40 |
| Temporal Mid | R | 3.97 | 58 | -14 | -16 |
| Postcentral | R | 3.97 | 50 | -14 | 32 |
| Cuneus | R | 3.94 | 2 | -78 | 32 |
| | R | 3.93 | 42 | -22 | 24 |
| Precuneus | R | 3.91 | 10 | -62 | 32 |
| | R | 3.91 | 34 | -14 | 16 |
| Precuneus | R | 3.9 | 10 | -54 | 40 |
| | R | 3.88 | 50 | -22 | 24 |
| Precuneus | L | 3.86 | -6 | -54 | 40 |
| Postcentral | R | 3.86 | 66 | -6 | 24 |
| | R | 3.86 | 42 | -22 | 32 |
| Postcentral | R | 3.82 | 58 | -22 | 32 |
| Cingulum Post | R | 3.82 | 10 | -54 | 32 |
| Postcentral | R | 3.81 | 66 | -14 | 32 |
| Cingulum Mid | L | 3.77 | -6 | -46 | 32 |
| Cingulum Mid | R | 3.77 | 2 | -46 | 32 |
| Calcarine | L | 3.77 | -6 | -62 | 16 |
| Calcarine | R | 3.76 | 10 | -70 | 16 |
| Postcentral | R | 3.76 | 66 | -14 | 16 |
| | R | 3.73 | 42 | -14 | 32 |
| Precentral | R | 3.73 | 50 | -6 | 24 |
| | R | 3.72 | 42 | -6 | 24 |
| Precuneus | R | 3.7 | 10 | -62 | 24 |
| Precuneus | R | 3.7 | 2 | -62 | 40 |
| Calcarine | R | 3.69 | 2 | -78 | 16 |
| Calcarine | R | 3.68 | 10 | -78 | 16 |
| Precuneus | L | 3.68 | -6 | -46 | 40 |
| Precuneus | R | 3.68 | 2 | -46 | 40 |
| | R | 3.66 | 50 | -30 | -8 |
| Temporal Sup | R | 3.65 | 50 | -14 | -8 |
| Postcentral | R | 3.65 | 58 | -14 | 24 |
| | R | 3.64 | 42 | -46 | 0 |
| | R | 3.64 | 42 | -38 | 0 |
| Precuneus | R | 3.64 | 10 | -54 | 24 |
| | R | 3.63 | 34 | -22 | -8 |

| | | | | | |
|---|---|---|---|---|---|
| | R | 3.63 | 50 | -46 | 0 |
| Heschl | R | 3.62 | 50 | -14 | 8 |
| | R | 3.62 | 34 | -38 | 8 |
| Precuneus | R | 3.61 | 2 | -54 | 48 |
| | R | 3.61 | 42 | -22 | -8 |
| Calcarine | L | 3.6 | -6 | -70 | 16 |
| | R | 3.59 | 26 | -22 | 16 |
| Temporal Mid | R | 3.56 | 66 | -14 | -16 |
| | R | 3.55 | 42 | -46 | -8 |
| Hippocampus | R | 3.55 | 26 | -38 | 8 |
| | L | 3.54 | -14 | -54 | 40 |
| Insula | R | 3.54 | 50 | -6 | 0 |
| Cuneus | L | 3.53 | -6 | -78 | 24 |
| Temporal Sup | R | 3.53 | 58 | -22 | 8 |
| Parietal Inf | R | 3.52 | 34 | -54 | 48 |
| Precuneus | L | 3.52 | -6 | -70 | 40 |
| Parietal Inf | R | 3.5 | 58 | -22 | 40 |
| | R | 3.49 | 34 | -6 | 16 |
| Rolandic Oper | R | 3.48 | 50 | 2 | 16 |
| Precuneus | R | 3.48 | 10 | -62 | 40 |
| Thalamus | R | 3.48 | 18 | -22 | 0 |
| | R | 3.46 | 26 | -30 | 16 |
| Precuneus | L | 3.46 | -6 | -62 | 48 |
| | R | 3.46 | 34 | -46 | 0 |
| Fusiform | R | 3.45 | 34 | -46 | -8 |
| | R | 3.45 | 34 | -14 | 24 |
| Precuneus | R | 3.45 | 2 | -70 | 32 |
| SupraMarginal | R | 3.45 | 58 | -22 | 16 |
| Cuneus | R | 3.44 | 2 | -86 | 24 |
| SupraMarginal | R | 3.43 | 58 | -22 | 24 |
| Insula | R | 3.43 | 42 | -6 | 0 |
| ParaHippocamp | R | 3.43 | 26 | -38 | -8 |
| | L | 3.41 | -14 | -54 | 32 |
| Cuneus | L | 3.41 | -14 | -62 | 32 |
| Precuneus | R | 3.4 | 18 | -62 | 32 |
| Temporal Mid | R | 3.4 | 58 | -14 | -24 |
| | R | 3.39 | 2 | -54 | 8 |
| Postcentral | R | 3.39 | 42 | -14 | 40 |
| | R | 3.38 | 34 | -14 | 32 |
| Precuneus | L | 3.37 | -6 | -54 | 48 |
| Postcentral | R | 3.37 | 50 | -22 | 40 |
| Calcarine | R | 3.36 | 10 | -62 | 8 |
| Occipital Sup | L | 3.36 | -22 | -62 | 32 |
| Hippocampus | R | 3.36 | 18 | -38 | 8 |
| Cingulum Post | R | 3.36 | 2 | -46 | 24 |
| Temporal Sup | R | 3.35 | 58 | -14 | 0 |
| Precuneus | R | 3.35 | 10 | -62 | 16 |
| Precentral | R | 3.33 | 58 | 2 | 16 |
| Precuneus | R | 3.33 | 10 | -46 | 40 |
| Precuneus | R | 3.33 | 10 | -54 | 48 |
| Cuneus | R | 3.31 | 10 | -70 | 24 |
| | R | 3.31 | 2 | -62 | 8 |

| | | | | | |
|---|---|---|---|---|---|
| Temporal Mid | R | 3.29 | 50 | -22 | -16 |
| | R | 3.29 | 42 | -54 | -8 |
| | R | 3.27 | 34 | -30 | 16 |
| | R | 3.27 | 42 | -38 | 8 |
| Precuneus | R | 3.27 | 18 | -54 | 40 |
| | R | 3.26 | 26 | -14 | 16 |
| Parietal Sup | L | 3.26 | -14 | -62 | 48 |
| Cingulum Post | R | 3.25 | 10 | -46 | 32 |
| Postcentral | R | 3.24 | 66 | -6 | 32 |
| Temporal Sup | R | 3.24 | 50 | -30 | 8 |
| Calcarine | R | 3.24 | 18 | -62 | 8 |
| Temporal Mid | R | 3.23 | 58 | -14 | -8 |
| Temporal Sup | R | 3.22 | 58 | -6 | 0 |
| Cuneus | L | 3.22 | -6 | -78 | 32 |
| SupraMarginal | R | 3.22 | 66 | -22 | 32 |
| Precuneus | R | 3.21 | 10 | -70 | 32 |
| Rolandic Oper | R | 3.2 | 42 | 2 | 16 |
| Precentral | R | 3.2 | 58 | 2 | 24 |
| Hippocampus | R | 3.2 | 18 | -30 | -8 |
| Occipital Mid | R | 3.19 | 26 | -62 | 40 |
| | R | 3.19 | 2 | -22 | 8 |
| Parietal Inf | R | 3.18 | 42 | -54 | 48 |
| | R | 3.18 | 34 | -22 | 24 |
| Putamen | R | 3.18 | 34 | -14 | 0 |
| Temporal Inf | R | 3.18 | 50 | -54 | -8 |
| | R | 3.17 | 26 | -46 | 0 |
| SupraMarginal | R | 3.16 | 50 | -30 | 24 |
| | R | 3.14 | 2 | -14 | 16 |
| Occipital Sup | L | 3.14 | -22 | -62 | 40 |
| Cingulum Post | R | 3.14 | 10 | -46 | 24 |
| | R | 3.13 | 26 | -54 | 40 |
| | R | 3.13 | 42 | -14 | -8 |
| Cuneus | L | 3.13 | -14 | -70 | 24 |
| Calcarine | L | 3.12 | -6 | -78 | 16 |
| Cuneus | R | 3.12 | 10 | -86 | 24 |
| Postcentral | R | 3.12 | 50 | -14 | 40 |
| Thalamus | R | 3.11 | 18 | -22 | 8 |
| | R | 3.11 | 50 | -22 | -8 |
| Cuneus | R | 3.1 | 2 | -86 | 32 |
| Precentral | R | 3.1 | 50 | -6 | 32 |
| Precentral | R | 3.1 | 50 | 2 | 24 |
| Precentral | R | 3.1 | 58 | 2 | 32 |
| Hippocampus | R | 3.08 | 26 | -30 | -8 |
| Putamen | R | 3.08 | 34 | -6 | 8 |
| | R | 3.08 | 18 | -54 | 32 |
| Rolandic Oper | R | 3.08 | 42 | -30 | 16 |
| Fusiform | R | 3.07 | 34 | -54 | -8 |
| Temporal Sup | R | 3.07 | 58 | -6 | -8 |
| Temporal Inf | R | 3.07 | 42 | -38 | -16 |
| Thalamus | R | 3.06 | 2 | -14 | 8 |
| Calcarine | R | 3.05 | 18 | -54 | 8 |
| Thalamus | L | 3.05 | -6 | -14 | 16 |

| | | | | | |
|---|---|---|---|---|---|
| | R | 3.04 | 26 | -46 | 40 |
| Cuneus | R | 3.04 | 10 | -86 | 16 |
| Precuneus | R | 3.04 | 10 | -54 | 16 |
| Precuneus | R | 3.04 | 18 | -62 | 40 |
| Thalamus | R | 3.04 | 10 | -30 | 8 |
| Insula | R | 3.03 | 42 | 2 | 8 |
| | R | 3.03 | 34 | -6 | 24 |
| Precuneus | R | 3.01 | 18 | -54 | 16 |
| Precuneus | L | 3.01 | -6 | -46 | 48 |
| | R | 3 | 2 | -46 | 16 |
| Rolandic Oper | R | 3 | 50 | 2 | 8 |
| Precuneus | L | 3 | -14 | -70 | 48 |
| Cingulum Post | R | 3 | 10 | -46 | 16 |

**Time window 10, cluster, infant vs. adult cry**

| Brain Region | Hemisphere | t-value | MNI Coordinates | | |
|---|---|---|---|---|---|
| | | | *x* | *y* | *z* |
| Precuneus | R | 6.36 | 2 | -62 | 32 |
| Cingulum Mid | R | 6.06 | 2 | -54 | 32 |
| Precuneus | L | 6.05 | -6 | -54 | 32 |
| Postcentral | R | 5.91 | 58 | -14 | 32 |
| Precuneus | L | 5.89 | -6 | -62 | 32 |
| Precuneus | R | 5.83 | 2 | -54 | 24 |
| Precuneus | R | 5.8 | 2 | -62 | 24 |
| Precuneus | L | 5.72 | -6 | -70 | 32 |
| Precuneus | L | 5.66 | -6 | -62 | 40 |
| Postcentral | R | 5.61 | 58 | -22 | 32 |
| Postcentral | R | 5.6 | 66 | -14 | 32 |
| Precuneus | R | 5.57 | 2 | -70 | 32 |
| Precuneus | L | 5.57 | -6 | -54 | 24 |
| Precuneus | R | 5.49 | 2 | -54 | 40 |
| Precentral | R | 5.46 | 58 | -6 | 32 |
| Precuneus | R | 5.4 | 2 | -62 | 40 |
| SupraMarginal | R | 5.18 | 66 | -22 | 32 |
| Heschl | R | 5.16 | 42 | -22 | 8 |
| Cuneus | R | 5.16 | 2 | -70 | 24 |
| Temporal Sup | R | 5.16 | 50 | -22 | 8 |
| Parietal Inf | R | 5.15 | 58 | -22 | 40 |
| Rolandic Oper | R | 5.14 | 50 | -22 | 16 |
| | R | 5.12 | 34 | -30 | 0 |
| SupraMarginal | R | 5.11 | 58 | -22 | 24 |
| Precuneus | L | 5.08 | -6 | -70 | 40 |
| | L | 5.06 | -14 | -62 | 40 |
| Precuneus | L | 5.06 | -6 | -62 | 24 |
| Precuneus | L | 5.05 | -6 | -54 | 40 |
| Heschl | R | 5.04 | 50 | -14 | 8 |
| Rolandic Oper | R | 5.01 | 42 | -22 | 16 |
| Temporal Sup | R | 5 | 58 | -22 | 8 |
| | R | 4.99 | 50 | -22 | 24 |
| Rolandic Oper | R | 4.98 | 50 | -14 | 16 |
| Cingulum Mid | R | 4.94 | 2 | -46 | 32 |
| Precentral | R | 4.94 | 50 | -6 | 32 |
| | R | 4.92 | 50 | -14 | 24 |
| Cingulum Post | R | 4.92 | 10 | -54 | 32 |
| SupraMarginal | R | 4.9 | 58 | -22 | 16 |
| Insula | R | 4.85 | 42 | -14 | 16 |
| Insula | R | 4.84 | 42 | -14 | 8 |
| Postcentral | R | 4.84 | 58 | -14 | 16 |
| Cingulum Post | R | 4.82 | 2 | -46 | 24 |
| Postcentral | R | 4.81 | 58 | -14 | 24 |
| Precuneus | R | 4.8 | 10 | -62 | 32 |
| Postcentral | R | 4.78 | 66 | -22 | 24 |
| Postcentral | R | 4.78 | 58 | -14 | 40 |
| Postcentral | R | 4.77 | 50 | -14 | 32 |
| Cingulum Mid | L | 4.75 | -6 | -46 | 32 |
| Precuneus | R | 4.74 | 10 | -54 | 40 |

| | | | | | |
|---|---|---|---|---|---|
| Postcentral | R | 4.73 | 50 | -22 | 32 |
| | R | 4.68 | 42 | -38 | -8 |
| Precentral | R | 4.68 | 50 | -6 | 24 |
| Precuneus | R | 4.68 | 10 | -62 | 24 |
| | L | 4.68 | -14 | -54 | 32 |
| Cuneus | L | 4.67 | -6 | -70 | 24 |
| Cuneus | L | 4.66 | -14 | -70 | 32 |
| | R | 4.66 | 26 | -30 | 0 |
| Cuneus | L | 4.64 | -14 | -62 | 32 |
| Hippocampus | R | 4.64 | 34 | -30 | -8 |
| Calcarine | R | 4.63 | 2 | -62 | 16 |
| Temporal Sup | R | 4.62 | 66 | -22 | 8 |
| Postcentral | R | 4.59 | 66 | -14 | 24 |
| Precuneus | R | 4.58 | 2 | -46 | 40 |
| Postcentral | R | 4.57 | 66 | -6 | 32 |
| | R | 4.55 | 42 | -22 | 0 |
| | R | 4.53 | 34 | -22 | 0 |
| Heschl | R | 4.51 | 42 | -30 | 8 |
| Temporal Sup | R | 4.51 | 50 | -22 | 0 |
| ParaHippocamp | R | 4.51 | 34 | -38 | -8 |
| | R | 4.48 | 2 | -54 | 16 |
| Precuneus | R | 4.48 | 10 | -46 | 40 |
| | R | 4.47 | 34 | -22 | 8 |
| Cuneus | R | 4.45 | 10 | -70 | 24 |
| Hippocampus | R | 4.42 | 34 | -38 | 0 |
| | R | 4.41 | 34 | -30 | 8 |
| Precuneus | L | 4.4 | -14 | -70 | 40 |
| Postcentral | R | 4.38 | 66 | -14 | 16 |
| | R | 4.38 | 42 | -22 | 24 |
| Precuneus | R | 4.38 | 2 | -54 | 48 |
| SupraMarginal | R | 4.36 | 66 | -22 | 16 |
| Cuneus | L | 4.35 | -6 | -78 | 40 |
| | R | 4.32 | 26 | -30 | 8 |
| Precuneus | L | 4.3 | -6 | -46 | 40 |
| | R | 4.3 | 26 | -46 | 48 |
| SupraMarginal | R | 4.29 | 66 | -22 | 40 |
| Insula | R | 4.29 | 42 | -14 | 0 |
| Cuneus | L | 4.29 | -6 | -78 | 32 |
| | R | 4.26 | 42 | -46 | -8 |
| Temporal Inf | R | 4.26 | 42 | -38 | -16 |
| Precuneus | R | 4.22 | 10 | -46 | 48 |
| Precuneus | L | 4.21 | -6 | -54 | 48 |
| Postcentral | R | 4.21 | 58 | -6 | 24 |
| Calcarine | L | 4.2 | -6 | -62 | 16 |
| | R | 4.2 | 42 | -14 | 24 |
| | L | 4.19 | -14 | -54 | 40 |
| Temporal Mid | R | 4.18 | 58 | -22 | 0 |
| Precuneus | R | 4.18 | 10 | -54 | 24 |
| | R | 4.17 | 50 | -46 | 0 |
| Precuneus | R | 4.14 | 10 | -54 | 48 |
| Precuneus | L | 4.13 | -6 | -62 | 48 |
| | R | 4.13 | 42 | -30 | 0 |

| | | | | | |
|---|---|---|---|---|---|
| Precuneus | R | 4.12 | 18 | -46 | 48 |
| Temporal Sup | R | 4.12 | 50 | -30 | 8 |
| Postcentral | R | 4.1 | 50 | -22 | 40 |
| | R | 4.06 | 42 | -6 | 32 |
| | R | 4.06 | 34 | -22 | 16 |
| Precentral | R | 4.04 | 58 | 2 | 40 |
| Calcarine | R | 4.04 | 10 | -62 | 8 |
| | R | 4.03 | 42 | -6 | 24 |
| Postcentral | R | 4.03 | 50 | -14 | 40 |
| Hippocampus | R | 4 | 26 | -38 | 0 |
| Postcentral | R | 3.99 | 58 | -6 | 40 |
| Thalamus | R | 3.98 | 26 | -22 | 0 |
| Heschl | R | 3.97 | 58 | -14 | 8 |
| Temporal Sup | R | 3.95 | 58 | -30 | 16 |
| Temporal Sup | R | 3.95 | 50 | -14 | 0 |
| Temporal Sup | R | 3.94 | 50 | -30 | 16 |
| Precuneus | R | 3.93 | 10 | -62 | 40 |
| Precuneus | R | 3.93 | 2 | -70 | 40 |
| | R | 3.92 | 50 | -46 | -8 |
| Postcentral | R | 3.92 | 34 | -38 | 48 |
| | R | 3.92 | 10 | -30 | 0 |
| | R | 3.9 | 42 | -46 | 0 |
| SupraMarginal | R | 3.9 | 58 | -30 | 24 |
| Precuneus | R | 3.9 | 10 | -62 | 16 |
| Rolandic Oper | R | 3.89 | 42 | -6 | 16 |
| | R | 3.89 | 42 | -22 | 32 |
| Cingulum Post | L | 3.88 | -6 | -46 | 24 |
| Temporal Mid | R | 3.87 | 50 | -54 | 0 |
| Rolandic Oper | R | 3.87 | 50 | -6 | 16 |
| Precentral | R | 3.86 | 50 | -6 | 40 |
| Precuneus | R | 3.85 | 2 | -46 | 48 |
| Rolandic Oper | R | 3.85 | 42 | -30 | 16 |
| Parietal Inf | R | 3.85 | 34 | -54 | 48 |
| | R | 3.84 | 26 | -22 | 8 |
| Cingulum Post | R | 3.81 | 10 | -46 | 32 |
| Putamen | R | 3.8 | 34 | -14 | 8 |
| | R | 3.79 | 42 | -14 | 32 |
| Parietal Inf | R | 3.78 | 34 | -46 | 48 |
| | R | 3.77 | 42 | -54 | 0 |
| | R | 3.76 | 42 | -30 | -8 |
| Precuneus | L | 3.75 | -6 | -54 | 16 |
| Thalamus | R | 3.75 | 18 | -30 | 0 |
| Precuneus | R | 3.73 | 10 | -70 | 32 |
| Fusiform | R | 3.72 | 34 | -46 | -8 |
| | R | 3.72 | 26 | -62 | 0 |
| Temporal Sup | R | 3.72 | 58 | -30 | 8 |
| Temporal Inf | R | 3.71 | 42 | -46 | -16 |
| Precuneus | R | 3.71 | 18 | -46 | 56 |
| SupraMarginal | R | 3.7 | 50 | -30 | 24 |
| | R | 3.69 | 2 | -46 | 16 |
| Precuneus | R | 3.69 | 2 | -62 | 48 |
| Parietal Sup | L | 3.68 | -14 | -62 | 48 |

| | | | | | |
|---|---|---|---|---|---|
| Temporal Inf | R | 3.67 | 50 | -54 | -8 |
| Precuneus | L | 3.67 | -6 | -70 | 48 |
| Temporal Sup | R | 3.67 | 66 | -14 | 8 |
| | R | 3.66 | 42 | -54 | -8 |
| Precuneus | R | 3.66 | 18 | -46 | 40 |
| Precuneus | R | 3.65 | 18 | -54 | 40 |
| Lingual | R | 3.63 | 18 | -62 | 0 |
| Calcarine | R | 3.63 | 18 | -62 | 8 |
| | R | 3.63 | 42 | -38 | 0 |
| Cuneus | L | 3.63 | -14 | -78 | 40 |
| Thalamus | R | 3.63 | 18 | -30 | 8 |
| Calcarine | R | 3.62 | 2 | -70 | 16 |
| Cuneus | R | 3.62 | 2 | -78 | 32 |
| Calcarine | L | 3.61 | -6 | -70 | 16 |
| Postcentral | R | 3.59 | 58 | -6 | 16 |
| Precentral | R | 3.59 | 42 | 2 | 32 |
| Cuneus | R | 3.59 | 2 | -78 | 24 |
| Postcentral | R | 3.58 | 66 | -6 | 24 |
| Temporal Mid | R | 3.58 | 50 | -30 | 0 |
| Cuneus | R | 3.57 | 10 | -78 | 24 |
| Temporal Mid | R | 3.56 | 58 | -54 | 0 |
| Temporal Sup | R | 3.56 | 66 | -30 | 16 |
| Temporal Inf | R | 3.56 | 42 | -30 | -16 |
| Precuneus | L | 3.55 | -14 | -62 | 24 |
| Cingulum Mid | R | 3.54 | 2 | -38 | 32 |
| Cuneus | L | 3.53 | -6 | -78 | 24 |
| | R | 3.52 | 50 | -22 | -8 |
| Hippocampus | R | 3.51 | 26 | -30 | -8 |
| Calcarine | R | 3.51 | 10 | -70 | 16 |
| Rolandic Oper | R | 3.5 | 50 | -6 | 8 |
| | L | 3.5 | -14 | -54 | 48 |
| | R | 3.48 | 26 | -54 | 40 |
| | R | 3.48 | 34 | -46 | 0 |
| Precuneus | R | 3.48 | 10 | -54 | 8 |
| | L | 3.46 | -14 | -46 | 40 |
| | R | 3.45 | 34 | -14 | 16 |
| ParaHippocamp | R | 3.44 | 26 | -38 | -8 |
| | R | 3.43 | 34 | -54 | 0 |
| Precuneus | R | 3.43 | 10 | -54 | 56 |
| | R | 3.43 | 50 | -30 | -8 |
| Precuneus | R | 3.43 | 2 | -70 | 48 |
| Insula | R | 3.43 | 42 | -6 | 8 |
| | R | 3.43 | 2 | -54 | 8 |
| Cingulum Mid | R | 3.43 | 2 | -38 | 40 |
| Postcentral | R | 3.42 | 42 | -22 | 40 |
| | R | 3.42 | 2 | -62 | 8 |
| Rolandic Oper | R | 3.42 | 58 | -6 | 8 |
| Postcentral | R | 3.42 | 42 | -14 | 40 |
| Parietal Sup | R | 3.4 | 34 | -46 | 56 |
| Parietal Inf | R | 3.4 | 58 | -22 | 48 |
| Cingulum Mid | L | 3.4 | -6 | -38 | 32 |
| Precuneus | R | 3.39 | 2 | -54 | 56 |

| | | | | | |
|---|---|---|---|---|---|
| Occipital Sup | L | 3.39 | -22 | -62 | 40 |
| Temporal Sup | R | 3.39 | 66 | -30 | 8 |
| Lingual | R | 3.39 | 10 | -62 | 0 |
| Temporal Inf | R | 3.38 | 50 | -38 | -16 |
| | R | 3.38 | 18 | -54 | 32 |
| Parietal Inf | R | 3.38 | 42 | -38 | 48 |
| Precuneus | R | 3.37 | 10 | -46 | 56 |
| Postcentral | R | 3.35 | 26 | -46 | 56 |
| Precuneus | L | 3.35 | -6 | -46 | 48 |
| Fusiform | R | 3.34 | 34 | -38 | -16 |
| Precuneus | L | 3.34 | -6 | -46 | 16 |
| | R | 3.34 | 34 | -62 | 0 |
| | R | 3.34 | 34 | -30 | 16 |
| | L | 3.32 | -14 | -46 | 32 |
| Precuneus | L | 3.32 | -14 | -70 | 48 |
| | R | 3.32 | 26 | -46 | 40 |
| Fusiform | R | 3.32 | 26 | -38 | -16 |
| | R | 3.32 | 34 | -14 | 32 |
| Hippocampus | R | 3.31 | 18 | -30 | -8 |
| Cuneus | L | 3.31 | -14 | -78 | 32 |
| Temporal Sup | R | 3.3 | 58 | -14 | 0 |
| Parietal Inf | R | 3.29 | 42 | -54 | 48 |
| | R | 3.28 | 34 | -6 | 32 |
| | R | 3.28 | 34 | -22 | 24 |
| Occipital Sup | L | 3.27 | -22 | -70 | 40 |
| Precuneus | R | 3.27 | 2 | -78 | 40 |
| Temporal Inf | R | 3.26 | 50 | -30 | -16 |
| | R | 3.26 | 42 | -38 | 8 |
| Postcentral | R | 3.26 | 66 | -6 | 16 |
| Rolandic Oper | R | 3.26 | 66 | -6 | 8 |
| Temporal Mid | R | 3.25 | 58 | -46 | -8 |
| Temporal Inf | R | 3.25 | 50 | -46 | -16 |
| Precentral | R | 3.25 | 42 | -6 | 40 |
| | R | 3.25 | 50 | -38 | -8 |
| Precuneus | R | 3.25 | 18 | -54 | 56 |
| Cingulum Mid | R | 3.24 | 2 | -38 | 48 |
| Cuneus | L | 3.24 | -14 | -70 | 24 |
| | R | 3.23 | 34 | -14 | 40 |
| Precuneus | R | 3.23 | 2 | -46 | 56 |
| Fusiform | R | 3.22 | 34 | -54 | -8 |
| Fusiform | R | 3.22 | 42 | -54 | -16 |
| Occipital Mid | R | 3.22 | 42 | -62 | 0 |
| Temporal Inf | R | 3.21 | 50 | -54 | -16 |
| | R | 3.21 | 34 | -38 | 8 |
| Precuneus | R | 3.21 | 18 | -54 | 48 |
| | L | 3.2 | -6 | -38 | 24 |
| Precuneus | L | 3.2 | -14 | -54 | 24 |
| Fusiform | R | 3.19 | 50 | -62 | -16 |
| Precentral | R | 3.19 | 50 | 2 | 32 |
| | R | 3.18 | 26 | -38 | 48 |
| | R | 3.18 | 42 | -22 | -8 |
| | R | 3.17 | 34 | -22 | -8 |

| | | | | | |
|---|---|---|---|---|---|
| Temporal Inf | R | 3.16 | 50 | -62 | -8 |
| Parietal Inf | R | 3.16 | 42 | -54 | 56 |
| | R | 3.16 | 10 | -30 | -8 |
| Precuneus | L | 3.15 | -6 | -78 | 48 |
| Cingulum Mid | L | 3.14 | -6 | -38 | 40 |
| Calcarine | R | 3.14 | 2 | -78 | 16 |
| Postcentral | R | 3.14 | 58 | -14 | 48 |
| Precuneus | R | 3.13 | 2 | -62 | 56 |
| Parietal Inf | R | 3.13 | 50 | -54 | 56 |
| Occipital Sup | L | 3.13 | -22 | -62 | 32 |
| SupraMarginal | R | 3.12 | 50 | -30 | 32 |
| Parietal Sup | R | 3.11 | 34 | -62 | 48 |
| Cingulum Mid | R | 3.11 | 10 | -38 | 40 |
| Temporal Mid | R | 3.08 | 50 | -22 | -16 |
| | R | 3.08 | 26 | -54 | 0 |
| Lingual | R | 3.07 | 18 | -38 | 0 |
| Temporal Mid | R | 3.07 | 66 | -22 | 0 |
| SupraMarginal | R | 3.07 | 58 | -30 | 32 |
| Precentral | R | 3.07 | 50 | 2 | 40 |
| Postcentral | R | 3.07 | 42 | -38 | 56 |
| Lingual | R | 3.06 | 10 | -70 | 8 |
| | R | 3.06 | 34 | -14 | 24 |
| Temporal Inf | R | 3.06 | 50 | -38 | -24 |
| Cuneus | L | 3.06 | -6 | -86 | 40 |
| Cuneus | L | 3.05 | -6 | -86 | 32 |
| Parietal Inf | R | 3.05 | 42 | -62 | 48 |
| Precuneus | R | 3.04 | 18 | -38 | 56 |
| Temporal Mid | R | 3.04 | 58 | -30 | 0 |
| Fusiform | R | 3.04 | 34 | -46 | -16 |
| Fusiform | R | 3.03 | 34 | -62 | -8 |
| Parietal Inf | R | 3.03 | 42 | -46 | 48 |
| Cingulum Mid | R | 3.03 | 18 | -38 | 48 |
| Lingual | R | 3.02 | 26 | -62 | -8 |
| ParaHippocamp | R | 3.02 | 34 | -30 | -16 |
| | R | 3 | 34 | -22 | 40 |

### Table ST2

**Time window 2, cluster 1, bonding and baby cry**

| **Brain Region** | **Hemisphere** | **r** | **MNI Coordinates** | | |
|---|---|---|---|---|---|
| | | | ***x*** | ***y*** | ***z*** |
| Lingual | R | -0.4 | 18 | -38 | 0 |
| ParaHippocamp | R | -0.41 | 34 | -38 | -8 |
| | R | -0.41 | 2 | -38 | 8 |
| | R | -0.41 | 18 | -22 | -8 |
| | R | -0.41 | 26 | -22 | -8 |
| ParaHippocamp | R | -0.41 | 26 | -38 | -8 |
| Temporal Inf | R | -0.42 | 42 | -38 | -16 |
| | R | -0.42 | 10 | -38 | -8 |
| | R | -0.43 | 42 | -30 | -8 |
| | R | -0.43 | 26 | -30 | 0 |
| Temporal Inf | R | -0.44 | 50 | -30 | -16 |
| ParaHippocamp | R | -0.44 | 34 | -30 | -16 |
| Fusiform | R | -0.46 | 26 | -38 | -16 |
| Thalamus | R | -0.47 | 18 | -30 | 0 |
| Fusiform | R | -0.47 | 34 | -38 | -16 |
| Hippocampus | R | -0.47 | 34 | -30 | -8 |
| | R | -0.48 | 10 | -38 | 0 |
| Lingual | R | -0.49 | 18 | -38 | -8 |
| Hippocampus | R | -0.51 | 26 | -30 | -8 |
| Hippocampus | R | -0.51 | 18 | -30 | -8 |
| | R | -0.53 | 2 | -30 | 0 |
| | R | -0.54 | 10 | -30 | 0 |
| Temporal Inf | R | -0.55 | 42 | -30 | -16 |

**Time window 2, cluster 2, bonding and baby cry**

| Brain Regic | Hemisphere | r | MNI Coordinates | | |
|---|---|---|---|---|---|
| | | | *x* | *y* | *z* |
| Subgenual | R | -0.4 | 2 | 10 | -8 |
| | R | -0.4 | 34 | -14 | 40 |
| SupraMargi | R | -0.4 | 66 | -38 | 24 |
| Postcentral | R | -0.4 | 50 | -22 | 40 |
| | R | -0.41 | 18 | -22 | 40 |
| | R | -0.41 | 2 | 10 | 16 |
| | R | -0.41 | 26 | -6 | 32 |
| | L | -0.41 | -6 | 2 | -8 |
| Cingulum M | R | -0.41 | 10 | -22 | 48 |
| Caudate | L | -0.41 | -6 | 10 | 8 |
| | R | -0.41 | 18 | -22 | 32 |
| Postcentral | R | -0.41 | 42 | -14 | 40 |
| Temporal M | R | -0.41 | 66 | -38 | 8 |
| | R | -0.41 | 10 | 10 | 24 |
| Supp Moto | R | -0.42 | 10 | -6 | 56 |
| Temporal S | R | -0.42 | 50 | -30 | 16 |
| SupraMargi | R | -0.42 | 50 | -30 | 32 |
| Postcentral | R | -0.42 | 42 | -30 | 40 |
| | R | -0.42 | 26 | -14 | 32 |
| Caudate | L | -0.42 | -6 | 10 | 0 |
| Temporal S | R | -0.42 | 66 | -30 | 8 |
| Heschl | R | -0.42 | 42 | -30 | 8 |
| Paracentr L | R | -0.42 | 10 | -22 | 56 |
| | R | -0.42 | 18 | -14 | 48 |
| Temporal M | R | -0.43 | 66 | -38 | 0 |
| Caudate | L | -0.43 | -6 | 10 | -8 |
| SupraMargi | R | -0.43 | 58 | -30 | 24 |
| Cingulum M | R | -0.43 | 10 | 2 | 32 |
| | R | -0.43 | 26 | -14 | 40 |
| | R | -0.43 | 18 | -6 | 40 |
| Cingulum M | R | -0.43 | 10 | -6 | 40 |
| Postcentral | R | -0.43 | 42 | -22 | 40 |
| | R | -0.44 | 18 | -14 | 40 |
| | R | -0.44 | 34 | -22 | 40 |
| Cingulum M | R | -0.44 | 10 | 2 | 40 |
| Temporal S | R | -0.45 | 42 | -38 | 16 |
| | L | -0.45 | -6 | 2 | 0 |
| | R | -0.45 | 18 | -6 | 32 |
| Temporal S | R | -0.45 | 66 | -30 | 16 |
| Temporal S | R | -0.45 | 58 | -30 | 8 |
| | R | -0.46 | 18 | 2 | 32 |
| Temporal S | R | -0.46 | 50 | -30 | 8 |
| Temporal M | R | -0.48 | 58 | -30 | 0 |
| SupraMargi | R | -0.5 | 66 | -30 | 24 |
| | R | -0.5 | 26 | -22 | 40 |
| SupraMargi | R | -0.5 | 58 | -38 | 32 |
| | R | -0.51 | 42 | -38 | 8 |
| Temporal M | R | -0.52 | 58 | -38 | 8 |

| | | | | |
|---|---|---|---|---|
| Temporal S R | -0.52 | 50 | -38 | 16 |
| Temporal S R | -0.53 | 66 | -38 | 16 |
| Temporal S R | -0.54 | 58 | -38 | 16 |
| Temporal M R | -0.55 | 66 | -30 | 0 |
| Temporal S R | -0.61 | 58 | -30 | 16 |
| Temporal S R | -0.61 | 50 | -38 | 8 |

**Time window 3, cluster 1, bonding and baby cry**

| Brain Region | Hemispher | r | MNI Coordinates | | |
|---|---|---|---|---|---|
| | | | *x* | *y* | *z* |
| | R | -0.4 | 42 | -22 | -8 |
| Temporal Mid | R | -0.4 | 58 | -30 | -8 |
| Rolandic Oper | R | -0.4 | 42 | -30 | 16 |
| Temporal Inf | R | -0.4 | 42 | -22 | -24 |
| SupraMarginal | R | -0.41 | 58 | -30 | 24 |
| Fusiform | R | -0.41 | 34 | -30 | -24 |
| ParaHippocamp | R | -0.41 | 34 | -22 | -24 |
| | R | -0.41 | 26 | -22 | -8 |
| Temporal Mid | R | -0.42 | 66 | -30 | -8 |
| Thalamus | R | -0.42 | 18 | -30 | 0 |
| Temporal Mid | R | -0.43 | 58 | -38 | 0 |
| Fusiform | R | -0.43 | 42 | -38 | -24 |
| | R | -0.43 | 50 | -30 | -8 |
| Hippocampus | R | -0.43 | 26 | -30 | -8 |
| | R | -0.43 | 18 | -22 | -8 |
| Temporal Mid | R | -0.43 | 66 | -46 | 8 |
| Hippocampus | R | -0.44 | 26 | -22 | -16 |
| | R | -0.44 | 42 | -22 | -16 |
| ParaHippocamp | R | -0.44 | 26 | -30 | -16 |
| Hippocampus | R | -0.44 | 18 | -30 | -8 |
| Temporal Inf | R | -0.44 | 42 | -30 | -24 |
| SupraMarginal | R | -0.44 | 66 | -30 | 24 |
| Temporal Inf | R | -0.44 | 50 | -30 | -24 |
| Hippocampus | R | -0.44 | 34 | -22 | -16 |
| Temporal Sup | R | -0.45 | 50 | -30 | 16 |
| Temporal Sup | R | -0.45 | 66 | -38 | 16 |
| Temporal Inf | R | -0.45 | 58 | -30 | -24 |
| | R | -0.46 | 42 | -30 | 0 |
| Temporal Sup | R | -0.46 | 66 | -46 | 16 |
| | R | -0.46 | 10 | -30 | 0 |
| Temporal Mid | R | -0.47 | 66 | -38 | 0 |
| Temporal Inf | R | -0.47 | 50 | -30 | -16 |
| | R | -0.47 | 42 | -30 | -8 |
| Heschl | R | -0.47 | 42 | -30 | 8 |
| Temporal Sup | R | -0.47 | 58 | -30 | 16 |
| Temporal Sup | R | -0.48 | 66 | -30 | 16 |
| Temporal Sup | R | -0.48 | 66 | -30 | 8 |
| Temporal Mid | R | -0.48 | 50 | -30 | 0 |
| Temporal Sup | R | -0.48 | 58 | -38 | 16 |
| Temporal Mid | R | -0.49 | 66 | -38 | 8 |
| Temporal Sup | R | -0.49 | 50 | -38 | 16 |
| Temporal Sup | R | -0.49 | 50 | -38 | 8 |
| ParaHippocamp | R | -0.5 | 34 | -30 | -16 |
| Temporal Mid | R | -0.5 | 58 | -38 | 8 |
| Hippocampus | R | -0.5 | 34 | -30 | -8 |
| Temporal Sup | R | -0.51 | 58 | -30 | 8 |
| Temporal Sup | R | -0.51 | 50 | -30 | 8 |
| Temporal Inf | R | -0.53 | 42 | -30 | -16 |
| Temporal Mid | R | -0.57 | 58 | -30 | 0 |

| | | | | | |
|---|---|---|---|---|---|
| Temporal Mid | R | -0.57 | 66 | -30 | 0 |

**Time window 3, cluster 2, bonding and baby cry**

| **Brain Regic** | **Hemisphere** | **r** | **MNI Coordinates** | | |
|---|---|---|---|---|---|
| | | | ***x*** | ***y*** | ***z*** |
| Insula | R | -0.4 | 42 | 10 | 8 |
| Front Inf O | R | -0.41 | 50 | 10 | 8 |
| Front Inf O | R | -0.41 | 58 | 10 | 8 |
| Postcentral | R | -0.41 | 66 | 2 | 16 |
| Front Inf O | R | -0.41 | 50 | 18 | 32 |
| Front Inf Tr | R | -0.42 | 50 | 18 | 24 |
| Front Inf O | R | -0.42 | 50 | 10 | 16 |
| Front Inf Tr | R | -0.42 | 58 | 26 | 24 |
| Front Inf Tr | R | -0.43 | 58 | 26 | 16 |
| Front Inf O | R | -0.44 | 58 | 10 | 16 |
| Front Inf Tr | R | -0.45 | 50 | 18 | 16 |
| Front Inf Tr | R | -0.46 | 58 | 18 | 24 |
| Front Inf Tr | R | -0.46 | 58 | 18 | 16 |
| Front Mid | R | -0.47 | 50 | 18 | 40 |
| Front Inf Tr | R | -0.48 | 50 | 18 | 8 |

**Time window 4, cluster, bonding and baby cry**

| **Brain Regic** | **Hemispher** | **r** | **MNI Coordinates** | | |
|---|---|---|---|---|---|
| | | | ***x*** | ***y*** | ***z*** |
| Temporal M | R | -0.41 | 50 | -30 | 0 |
| Heschl | R | -0.41 | 42 | -30 | 8 |
| SupraMargi | R | -0.41 | 66 | -30 | 24 |
| Temporal S | R | -0.42 | 66 | -30 | 8 |
| Temporal M | R | -0.42 | 66 | -30 | 0 |
| Temporal S | R | -0.42 | 50 | -38 | 16 |
| Temporal S | R | -0.43 | 58 | -30 | 8 |
| Temporal S | R | -0.43 | 50 | -30 | 8 |
| Temporal M | R | -0.45 | 66 | -38 | 0 |
| Temporal S | R | -0.45 | 50 | -38 | 8 |
| Temporal M | R | -0.46 | 66 | -38 | 8 |
| Temporal S | R | -0.46 | 58 | -38 | 16 |
| Temporal M | R | -0.47 | 58 | -30 | 0 |
| Temporal S | R | -0.47 | 66 | -30 | 16 |
| Temporal M | R | -0.48 | 58 | -38 | 8 |
| Temporal S | R | -0.5 | 66 | -38 | 16 |

**Time window 5, cluster, bonding and baby cry**

| **Brain Region** | **Hemisphere** | **r** | **MNI Coordinates** | | |
|---|---|---|---|---|---|
| | | | ***x*** | ***y*** | ***z*** |
| Temporal Inf | R | -0.42 | 50 | -62 | -8 |
| Occipital Mid | R | -0.45 | 42 | -62 | 8 |
| Temporal Mid | R | -0.47 | 50 | -70 | 8 |
| Occipital Inf | R | -0.48 | 50 | -70 | -8 |
| Occipital Mid | R | -0.54 | 50 | -70 | 0 |

**Time window 6, cluster 1, bonding and baby cry**

| **Brain Region** | **Hemisphere** | **r** | **MNI Coordinates** | | |
|---|---|---|---|---|---|
| | | | ***x*** | ***y*** | ***z*** |
| Precuneus | R | -0.4 | 2 | -46 | 48 |
| Cingulum Mic | R | -0.41 | 2 | -38 | 32 |
| Cingulum Mic | R | -0.42 | 2 | -38 | 40 |
| Cingulum Pos | R | -0.42 | 10 | -38 | 32 |
| Cingulum Mic | R | -0.45 | 10 | -38 | 40 |

**Time window 6, cluster 2, bonding and baby cry**

| **Brain Region** | **Hemisphere** | **r** | **MNI Coordinates** | | |
|---|---|---|---|---|---|
| | | | ***x*** | ***y*** | ***z*** |
| Occipital Mid | R | -0.41 | 50 | -70 | 0 |
| | R | -0.42 | 42 | -62 | 16 |
| Occipital Mid | R | -0.42 | 42 | -62 | 0 |
| Temporal Mid | R | -0.42 | 50 | -70 | 8 |
| | R | -0.45 | 42 | -54 | 8 |
| Occipital Inf | R | -0.46 | 50 | -70 | -8 |
| Temporal Inf | R | -0.47 | 50 | -62 | -8 |
| Occipital Mid | R | -0.53 | 42 | -62 | 8 |

**Time window 7, cluster, bonding and baby cry**

| **Brain Region** | **Hemisphere** | **r** | **MNI Coordinates** | | |
|---|---|---|---|---|---|
| | | | ***x*** | ***y*** | ***z*** |
| Temporal Sup | R | -0.42 | 50 | -38 | 8 |
| | R | -0.45 | 34 | -38 | 8 |
| | R | -0.45 | 42 | -38 | 8 |
| Temporal Sup | R | -0.46 | 42 | -38 | 16 |

### Table ST3: actigraphy settings

| argument | value | context |
|---|---|---|
| GGIR_versio | Could not ret | not applicable |
| R_version | R version 3.6 | not applicable |
| backup.cal.cc | retrieve | Calibration, Feature extraction, Epoch size, Time zone |
| chunksize | 1 | Calibration, Feature extraction, Epoch size, Time zone |
| configtz | c() | Calibration, Feature extraction, Epoch size, Time zone |
| dayborder | 0 | Calibration, Feature extraction, Epoch size, Time zone |
| do.anglex | FALSE | Calibration, Feature extraction, Epoch size, Time zone |
| do.angley | FALSE | Calibration, Feature extraction, Epoch size, Time zone |
| do.anglez | TRUE | Calibration, Feature extraction, Epoch size, Time zone |
| do.bfen | FALSE | Calibration, Feature extraction, Epoch size, Time zone |
| do.cal | TRUE | Calibration, Feature extraction, Epoch size, Time zone |
| do.dev_roll_r | FALSE | Calibration, Feature extraction, Epoch size, Time zone |
| do.dev_roll_r | FALSE | Calibration, Feature extraction, Epoch size, Time zone |
| do.dev_roll_r | FALSE | Calibration, Feature extraction, Epoch size, Time zone |
| do.en | FALSE | Calibration, Feature extraction, Epoch size, Time zone |
| do.enmo | TRUE | Calibration, Feature extraction, Epoch size, Time zone |
| do.enmoa | FALSE | Calibration, Feature extraction, Epoch size, Time zone |
| do.hfen | FALSE | Calibration, Feature extraction, Epoch size, Time zone |
| do.hfenplus | FALSE | Calibration, Feature extraction, Epoch size, Time zone |
| do.lfen | FALSE | Calibration, Feature extraction, Epoch size, Time zone |
| do.lfenmo | FALSE | Calibration, Feature extraction, Epoch size, Time zone |
| do.mad | FALSE | Calibration, Feature extraction, Epoch size, Time zone |
| do.roll_med_ | FALSE | Calibration, Feature extraction, Epoch size, Time zone |
| do.roll_med_ | FALSE | Calibration, Feature extraction, Epoch size, Time zone |
| do.roll_med_ | FALSE | Calibration, Feature extraction, Epoch size, Time zone |
| dynrange | c() | Calibration, Feature extraction, Epoch size, Time zone |
| hb | 15 | Calibration, Feature extraction, Epoch size, Time zone |
| lb | 0.5 | Calibration, Feature extraction, Epoch size, Time zone |
| minloadcrit | 72 | Calibration, Feature extraction, Epoch size, Time zone |
| myfun | c() | Calibration, Feature extraction, Epoch size, Time zone |
| n | 4 | Calibration, Feature extraction, Epoch size, Time zone |
| print.filenam | FALSE | Calibration, Feature extraction, Epoch size, Time zone |
| printsummar | FALSE | Calibration, Feature extraction, Epoch size, Time zone |
| windowsizes | c(5,900,3600 | Calibration, Feature extraction, Epoch size, Time zone |
| acc.metric | ENMO | General parameters |
| bout.metric | 4 | General parameters |
| config_file_i | /projects/MI | General parameters |
| data_cleanin | c() | General parameters |
| datadir | /projects/MI | General parameters |
| desiredtz | | General parameters |
| do.bfx | FALSE | General parameters |
| do.bfy | FALSE | General parameters |
| do.bfz | FALSE | General parameters |
| do.hfx | FALSE | General parameters |
| do.hfy | FALSE | General parameters |
| do.hfz | FALSE | General parameters |

| | | |
|---|---|---|
| do.lfx | FALSE | General parameters |
| do.lfy | FALSE | General parameters |
| do.lfz | FALSE | General parameters |
| do.parallel | TRUE | General parameters |
| do.report | c(2,4,5) | General parameters |
| do.sgAccEN | TRUE | General parameters |
| do.sgAnglex | FALSE | General parameters |
| do.sgAngley | FALSE | General parameters |
| do.sgAnglez | FALSE | General parameters |
| excludefirst.p | FALSE | General parameters |
| excludelast.p | FALSE | General parameters |
| f0 | 1 | General parameters |
| f1 | 50 | General parameters |
| GGIRversion | 2.3-0 | General parameters |
| i | 30 | General parameters |
| idloc | 1 | General parameters |
| includedaycri | 0.66666667 | General parameters |
| minimum_M | 23 | General parameters |
| minimumFile | 2 | General parameters |
| mode | c(1,2,3,4,5) | General parameters |
| MX.ig.min.du | 10 | General parameters |
| outputdir | /projects/MI | General parameters |
| overwrite | FALSE | General parameters |
| part5_agg2_ | FALSE | General parameters |
| relyonguider | FALSE | General parameters |
| rmc.bitrate | c() | General parameters |
| rmc.check4ti | FALSE | General parameters |
| rmc.col.acc | c(1,2,3) | General parameters |
| rmc.col.temp | c() | General parameters |
| rmc.col.time | c() | General parameters |
| rmc.col.wear | c() | General parameters |
| rmc.dec | . | General parameters |
| rmc.desiredtz | | General parameters |
| rmc.doresam | FALSE | General parameters |
| rmc.dynamic | c() | General parameters |
| rmc.firstrow. | c() | General parameters |
| rmc.firstrow. | c() | General parameters |
| rmc.format.t | %Y-%m-%d % | General parameters |
| rmc.header.l | c() | General parameters |
| rmc.header.s | c() | General parameters |
| rmc.headern | c() | General parameters |
| rmc.headern | c() | General parameters |
| rmc.headern | c() | General parameters |
| rmc.noise | FALSE | General parameters |
| rmc.origin | 01/01/1970 | General parameters |
| rmc.sf | c() | General parameters |
| rmc.unit.acc | g | General parameters |
| rmc.unit.tem | C | General parameters |

| | | |
|---|---|---|
| rmc.unit.tim | POSIX | General parameters |
| rmc.unsigne | TRUE | General parameters |
| save_ms5rav | csv | General parameters |
| save_ms5rav | TRUE | General parameters |
| selectdaysfil | c() | General parameters |
| SI | c(list(platfo | |
| storefolderst | FALSE | General parameters |
| studyname | c() | General parameters |
| anglethreshc | 5 | Parameters sleep detection |
| constrain2ra | TRUE | Parameters sleep detection |
| ignorenonwe | TRUE | Parameters sleep detection |
| timethreshol | 5 | Parameters sleep detection |
| colid | 1 | Parameters sleep period time detection with or wihout sleeplog |
| coln1 | 2 | Parameters sleep period time detection with or wihout sleeplog |
| criterror | 4 | Parameters sleep period time detection with or wihout sleeplog |
| def.noc.sleep | c(21,8) | Parameters sleep period time detection with or wihout sleeplog |
| do.visual | TRUE | Parameters sleep period time detection with or wihout sleeplog |
| excludefirstla | FALSE | Parameters sleep period time detection with or wihout sleeplog |
| includenight | 16 | Parameters sleep period time detection with or wihout sleeplog |
| loglocation | /projects/MI | Parameters sleep period time detection with or wihout sleeplog |
| nnights | 7 | Parameters sleep period time detection with or wihout sleeplog |
| outliers.only | TRUE | Parameters sleep period time detection with or wihout sleeplog |
| relyonsleeplc | c() | Parameters sleep period time detection with or wihout sleeplog |
| sleeplogidnu | TRUE | Parameters sleep period time detection with or wihout sleeplog |
| boutcriter.in | 0.9 | Parameters time-use variables |
| boutcriter.lig | 0.8 | Parameters time-use variables |
| boutcriter.m | 0.8 | Parameters time-use variables |
| boutdur.in | c(1,10,30) | Parameters time-use variables |
| boutdur.lig | c(1,10) | Parameters time-use variables |
| boutdur.mvp | 1 | Parameters time-use variables |
| excludefirstla | FALSE | Parameters time-use variables |
| save_ms5rav | FALSE | Parameters time-use variables |
| threshold.lig | 30 | Parameters time-use variables |
| threshold.mc | 100 | Parameters time-use variables |
| threshold.vig | 400 | Parameters time-use variables |
| timewindow | MM | Parameters time-use variables |
| boutcriter | 0.8 | Study design, Parameters descriptive analysis |
| closedbout | FALSE | Study design, Parameters descriptive analysis |
| do.imp | TRUE | Study design, Parameters descriptive analysis |
| do.part3.pdf | TRUE | Study design, Parameters descriptive analysis |
| epochvalues2 | FALSE | Study design, Parameters descriptive analysis |
| hrs.del.end | 0 | Study design, Parameters descriptive analysis |
| hrs.del.start | 0 | Study design, Parameters descriptive analysis |
| iglevels | c() | Study design, Parameters descriptive analysis |
| ilevels | c() | Study design, Parameters descriptive analysis |
| includedaycri | 16 | Study design, Parameters descriptive analysis |
| IVIS_epochsi | c() | Study design, Parameters descriptive analysis |

| | | |
|---|---|---|
| IVIS_window | 60 | Study design, Parameters descriptive analysis |
| IVIS.activity.r | 2 | Study design, Parameters descriptive analysis |
| M5L5res | 10 | Study design, Parameters descriptive analysis |
| maxdur | 7 | Study design, Parameters descriptive analysis |
| mvpadur | c(1,5,10) | Study design, Parameters descriptive analysis |
| mvpathreshc | 100 | Study design, Parameters descriptive analysis |
| ndayswindov | 7 | Study design, Parameters descriptive analysis |
| qlevels | c() | Study design, Parameters descriptive analysis |
| qM5L5 | c() | Study design, Parameters descriptive analysis |
| qwindow | c(0,24) | Study design, Parameters descriptive analysis |
| strategy | 3 | Study design, Parameters descriptive analysis |
| TimeSegmer | c() | Study design, Parameters descriptive analysis |
| window.sum | 10 | Study design, Parameters descriptive analysis |
| winhr | 5 | Study design, Parameters descriptive analysis |
| dofirstpage | TRUE | Visual report |
| viewingwind | 1 | Visual report |
| visualreport | TRUE | Visual report |

The original excel sheet can be provided on reasonable request
